# An adenylyl cyclase switch reroutes carbon from growth to virulence lipids in *Mycobacterium tuberculosis*

**DOI:** 10.64898/2026.09.10.750769

**Authors:** Selva Rupa Christinal Immanuel, Julie Do, Eliza J. R. Peterson, Kristopher A. Hunt, Amardeep Kaur, Min Pan, Albel Singh, Wei-Ju Wu, Apoorva M. Bhatt, Nitin S. Baliga

**Affiliations:** Institute for Systems Biology, Seattle, WA 98109, USA; School of Biosciences and Institute of Microbiology and Infection, University of Birmingham, Birmingham B15 2TT, UK; Departments of Biology and Microbiology, University of Washington, Seattle, WA 98195, USA; Molecular and Cellular Biology Program, University of Washington, Seattle, WA 98195, USA; Lawrence Berkeley National Lab, Berkeley, CA 94720, USA

**Keywords:** (i) *Mycobacterium tuberculosis*, (ii) Rv1625c (adenylyl cyclase), (iii) cAMP signaling, (iv) electron transport chain, (v) central carbon metabolism, (vi) phthiocerol dimycocerosate (PDIM), (vii) genome-scale metabolic modeling, (viii) antitubercular drug mechanism of action, (ix) GSK 2556286 (GSK-286), (x) TBD11 (mCLB073)

## Abstract

Anti-TB drugs act non-uniformly on *Mycobacterium tuberculosis* (Mtb) because of its distinct physiological states across diverse infection niches shaped by host-derived nutrients and stresses. Agonists of the adenylyl cyclase Rv1625c are a novel drug class that selectively inhibits Mtb growth in macrophages and cholesterol-rich conditions by an unknown mechanism. Combining condition-resolved transcriptomics, genome-scale metabolic modeling, and genetic perturbation, we show these agonists cause a blockade in the electron transport chain, which likely activates Rv1625c. The elevated cAMP in turn drives global transcriptional and post-translational remodeling of central carbon and lipid metabolism. The resulting methylcitrate cycle reversal chokes cholesterol breakdown products from entering central metabolism, diverting carbon toward cell wall and virulence-lipid (phthiocerol dimycocerosate) synthesis and inhibiting growth. These drugs thus hijack an endogenous switch that reroutes carbon from biomass to virulence-lipid production. Hence, nutrients that restore carbon flux and relieve ETC blockade reduce activity, whereas Rv1625c overexpression and ETC inhibitors potentiate drug action even in refractory conditions.

## INTRODUCTION

*Mycobacterium tuberculosis* (Mtb), the causative agent of tuberculosis, persists in the host by adopting metabolically diverse and heterogeneous states. This phenotypic heterogeneity arises from the varied microenvironments encountered during infection, ranging from oxygen-rich to hypoxic niches and from nutrient-replete to nutrient-deprived conditions. Within these niches, Mtb flexibly remodels its metabolism to utilize host-derived carbon sources such as fatty acids, cholesterol, and short-chain organic acids including propionate, adaptations that underpin survival, replication, and immune evasion through tight regulation of carbon catabolism and energy generation^1–14^. This same flexibility has a profound therapeutic consequence: because Mtb’s physiologic state is dictated by the nutrients available in each niche, many anti-tubercular drugs act non-uniformly across infection contexts, confounding both their discovery and the interpretation of their mechanisms^3,15–20^.

Cholesterol metabolism is especially central to the host-adapted state, integrating catabolic and regulatory functions specific to the intracellular environment. Large gene clusters required for cholesterol uptake and steroid-ring degradation, such as *hsaA-hsaD*, *fadD3*, and *ipdAB*, are repressed by the transcription factor KstR^21–25^. Cholesterol-derived propionyl-CoA is detoxified through the methylcitrate cycle (MCC; *prpC*, *prpD*, *icl1*) and the glyoxylate shunt (*icl1*, *icl2*), with PrpR coordinating the *prpDC* operon^26,27^. Odd-chain fatty acids likewise generate propionyl-CoA, whereas even-chain fatty acids yield acetyl-CoA that feeds the TCA and glyoxylate cycles^3^. Beyond supplying carbon, cholesterol catabolism imposes a substantial reductive burden on the cell, generating large amounts of NADH and thereby directly linking lipid breakdown to the redox and respiratory state of the cell^28^.

This coupling of carbon catabolism to redox balance is a recurring theme in Mtb’s conditional physiology. Central carbon metabolism, the TCA cycle, and the electron transport chain (ETC) are extensively reprogrammed according to nutrient and oxygen availability, such that enzymes including isocitrate lyase (Icl1)^25,27–30^, fumarate reductase (Frd)^31^, and malate dehydrogenase (Mdh) or malate quinone oxidoreductase (Mqo)^32,33^ display context-dependent essentiality. Under hypoxia or reductive stress, Mtb engages the glyoxylate shunt or a partially reductive TCA cycle, with fumarate reductase (*frdABCD*) helping to maintain redox balance and persistence^31,34,35^. Substrate-dependent flux rewiring is similarly evident in the reversal of MCC flux required to recycle propionate during growth on pyruvate or lactate^36,37^, and in the strong dependence of glycerol-grown cells on cytochrome *bd* oxidase (*cydAB*) when the cytochrome *bc1-aa3* complex (*qcrCAB*, *ctaCDE*) is impaired^18^. Additional redox flexibility is provided by the type I (*nuoA-N*) and type II (*ndh*, *ndhA*) NADH dehydrogenases^38–46^. Layered on top of this metabolic network, the second-messenger cAMP serves as a major integrator of stress and metabolism; Mtb encodes an unusually large repertoire of adenylyl cyclases, several tuned to distinct environmental cues such as the pH-sensitive Rv1264^47–53^.

Among these cyclases, Rv1625c has emerged as a compelling drug target^53–59^. Rv1625c agonists were discovered by screening compounds against Mtb-infected macrophages, and their activity is recapitulated *in vitro* in media containing cholesterol, one of the preferred carbon sources of intracellular Mtb^56–59^. Small-molecule agonists such as V-58^57^ and GSK2556286 (GSK286)^56,58,59^ rapidly induce cAMP in an Rv1625c-dependent manner. Disruption of Rv1625c abolishes drug activity, whereas its overexpression potentiates killing, and resistant mutants map to the cyclase itself^58^. Rv1625c resembles an ancestral form of mammalian membrane adenylyl cyclases^53,60^ and is itself lipid-regulated, with its hexa-helical membrane anchor acting as an inhibitory receptor for fatty acids, particularly oleic acid^61^. Yet how these agonists actually arrest Mtb growth has remained unresolved. CO₂-release assays and fluorescent reporters of the cholesterol breakdown product propionyl-CoA showed that drug treatment blocks cholesterol catabolism, with no CO₂ evolved and no propionyl-CoA accumulation, leading to the proposal that the compounds prevent cholesterol uptake^58,59^. Critically, elevated cAMP alone neither inhibits growth nor explains the strict carbon-source dependence of drug activity^57–59^. Thus, beyond the requirement for Rv1625c and the observation that cholesterol catabolism is blocked, the downstream events that convert cAMP signaling into a growth-inhibitory, context-specific phenotype have been unknown, an explanatory gap that target-centric approaches are poorly equipped to close for a drug whose efficacy is set by the metabolic state of the cell.

We reasoned that resolving such a mechanism requires a systems-level view of the metabolic network the drug perturbs. Here we combined condition-resolved transcriptomics, an updated genome-scale metabolic model, flux balance analysis, and targeted genetic perturbation to define the action of two Rv1625c agonists – GSK286 and TBD11 (mCLB073), across host-relevant carbon sources. This approach revealed that the blockade in cholesterol metabolism is not a block in uptake but a consequence of a reversal of the MCC, driven by cAMP-mediated global remodeling of central carbon and lipid metabolism at both the transcriptional and post-translational levels. We find that this rewiring diverts carbon away from biomass and toward synthesis of cell wall lipids, including the virulence lipid phthiocerol dimycocerosate (PDIM). Further, our findings suggest that the agonists most likely act upstream by impairing the ETC, such that the resulting perturbed redox state is what potentially activates Rv1625c and sets in motion the cascade that accounts for the prior observations. Together, these findings establish the metabolic context of drug action as a central determinant of efficacy and uncover a fundamental mechanism by which Mtb toggles carbon allocation between virulence-lipid production and growth.

## RESULTS

### Rv1625c agonists induce CRP-mediated global transcriptome remodeling, even in contexts in which they do not cause growth-inhibition

Building on previously reported conditional activity of GSK286^56–59^, we first investigated whether the activity of TBD11 (mCLB073) was carbon source-dependent and required a functional copy of Rv1625c. We performed dose response assays to measure the minimum inhibitory concentration (**MIC**) of each of the two drugs on Mtb H37Rv with and without CRISPRi knockdown (**KD**) of Rv1625c in modified Sauton’s medium supplemented with different carbon sources. As expected, both drugs inhibited growth of Mtb in the presence of cholesterol or propionate (here onwards “**inhibitory conditions**”), but neither had any growth inhibitory effect on Mtb cultures in glycerol or cholesterol + acetate (here onwards “**non-inhibitory conditions**”). Knockdown of Rv1625c abolished activity of both GSK286 and TBD11 in inhibitory conditions, confirming the essential role of Rv1625c in the mechanism of action of these drugs (**Figure 1A and 1B**), which has been shown to be associated with elevated cAMP levels ^56–59^. Given the established role of cAMP as a mediator of global transcriptional regulation^52,62–67^, we performed comprehensive transcriptome profiling of Mtb following GSK286 treatment under both inhibitory and non-inhibitory conditions to uncover clues into the mechanism of carbon source-dependent growth inhibition by Rv1625c agonists. We profiled transcriptomes of Mtb cultures in modified Sauton’s medium with cholesterol or cholesterol + acetate over a time course, pre- and post-treatment with 5X (0.05 μg/mL) and 15X (1.5 μg/mL) GSK286. Altogether, 428 (264 up and 181 down) and 262 (195 up and 74 down) genes were differentially regulated over the time course of treatment in cultures with cholesterol and cholesterol + acetate, respectively. Despite the distinct phenotypic consequences, key genes of central carbon metabolism (**CCM**), including the TCA cycle and the anaplerotic pathway, exhibited remarkably similar patterns of down- and up-regulation across inhibitory and non-inhibitory conditions (**Figure 1C-E**).

**Figure 1:**
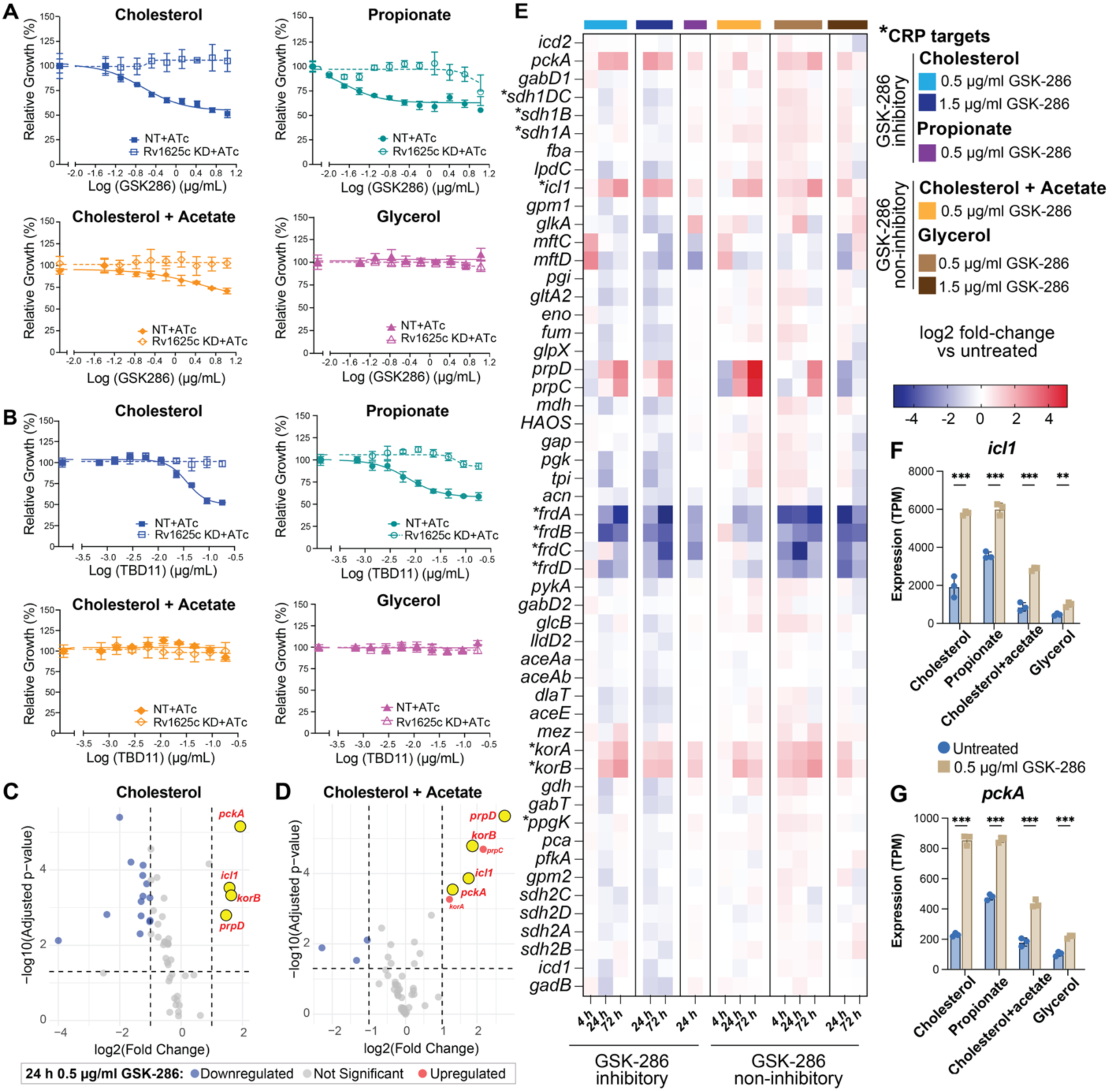
C-source-dependent growth inhibition of Mtb by Rv1625c agonists and the associated global differential gene regulation. Relative growth inhibition of Mtb by Rv1625c agonists, GSK286 (**A**) and TBD11 (**B**), in dose-response assays across “inhibitory” (cholesterol and propionate) and “non-inhibitory” (glycerol and cholesterol + acetate) conditions. Each panel includes non-targeting (NT) sgRNA control and CRISPRi knockdown (KD) of Rv1625c (dashed lines in all plots). Data are representative of at least two independent experiments. Error bars correspond to standard deviation of replicates within the same experiment. (**C-D**) Volcano plots show significant differential regulation of CCM genes across inhibitory and non-inhibitory conditions; key genes of the TCA cycle and the anaplerotic pathway are highlighted. (**E**) Differential regulation of CCM genes under inhibitory and non-inhibitory conditions (see inset key for specific growth contexts and color-scale for differential expression levels; asterisks indicate known targets of CRP). (**F-G**) Gene expression values (reported as Transcripts Per Million (TPM)) of key enzymes of the TCA cycle (*icl1*) and the anaplerotic pathway (*pckA*) under inhibitory and non-inhibitory conditions. P-values are deduced using two-way ANOVA, where **<0.01, ***<0.001, and ****<0.0001.

As expected, the DEGs were significantly enriched (p-value = 2.34x10⁻⁸) for 40 genes (20 upregulated and 20 downregulated) that are known regulatory targets of the cAMP receptor protein (CRP)^67^, based on experimentally-mapped CRP-binding sites in their promoters and their differential expression upon CRP deletion^67^. These findings provide strong evidence for CRP involvement in the response to Rv1625c agonists. Noteworthy CRP-regulated genes among the DEGs were *icl1* (required for cholesterol utilization)^25,27,29,30^ and *pckA* (required for cholesterol as well as glycerol metabolism)^68–70^, both significantly upregulated following GSK286 treatment across inhibitory and non-inhibitory conditions (**Figure 1F and 1G**). These findings demonstrate that while the increased cAMP levels upon induction of Rv1625c drives global CRP-mediated transcriptional remodeling, this phenomenon occurs across both inhibitory and non-inhibitory conditions. This finding was consistent with prior reports ^56–59^ that treatment with RV1625c-agonists resulted in elevated cAMP levels, even in non-inhibitory conditions. In other words, the carbon source-dependent growth inhibition by Rv1625c-agonists was not explained at the level of transcriptional regulation of CCM, motivating further inquiry into downstream consequences on metabolism.

### Global remodeling of metabolism by elevated cAMP levels explains context-dependent growth inhibition by Rv1625c-agonists

In order to elucidate how transcriptional changes cause context-dependent growth inhibition by Rv1625c agonists, we curated a genome-scale constraints-based metabolic model to incorporate recent advances in the understanding of Mtb metabolism (**Table 1**). Through an iterative process, we systematically curated 113 reactions of iEK1011^71^ and improved mechanistic accuracy of the model in recapitulating experimentally characterized growth on various C-substrates. The updated metabolic model (iSI1012) is comprised of 1,012 enzymes that catalyze the transformation of 974 metabolites through 1,232 reactions. Key model improvements included making unidirectional reactions (2MCD, 2MCS, 2MID, and 2MIL) bidirectional to allow the methyl citrate cycle (**MCC**) to operate in reverse during utilization of glycerol, glucose, pyruvate, and lactate^36,72^, while maintaining forward direction with cholesterol and acetate. Additional updates included removing the assignment of LldD1 (Rv0694) and leaving LldD2 (*Rv1872c*)^73^ as the sole essential lactate dehydrogenase responsible for conversion of lactate to pyruvate via the L-lactate dehydrogenase reaction (L_LACD)^37,73^, removal of Rv1127c assignment to the PPDK reaction^72,74^, curation of ETC reactions to revise quinone assignments, stoichiometry of electron transfer and proton translocation by non-proton pumping cytochrome BD oxidases^75–77^ and NADH dehydrogenases reactions^44–46,78,79^, and the incorporation of electron transfer flavoprotein reactions^80^ (see **Table 1** and **Data S1** for a full account of evidence-based curation of all 113 reactions). The curated model was then contextualized with transcriptome profiles to uncover metabolic states of Mtb with and without GSK286 treatment, across both inhibitory and non-inhibitory conditions. In addition to accurately recapitulating known baseline flux states of Mtb^81^ during growth on different carbon sources (**Table 1**, **Figure 2A**), flux balance analysis (**FBA**)^82–85^ with the GIMME^83^ contextualized models also recapitulated growth inhibition by drug treatment in inhibitory conditions, but no phenotypic consequence in non-inhibitory conditions^56–59^ (**Figure 2B; Data S2,** GitHub repository https://github.com/baliga-lab/iSI1012-metabolic-model-of-Mtb-and-Rv1625c-agonists-characterization).

**Figure 2.**
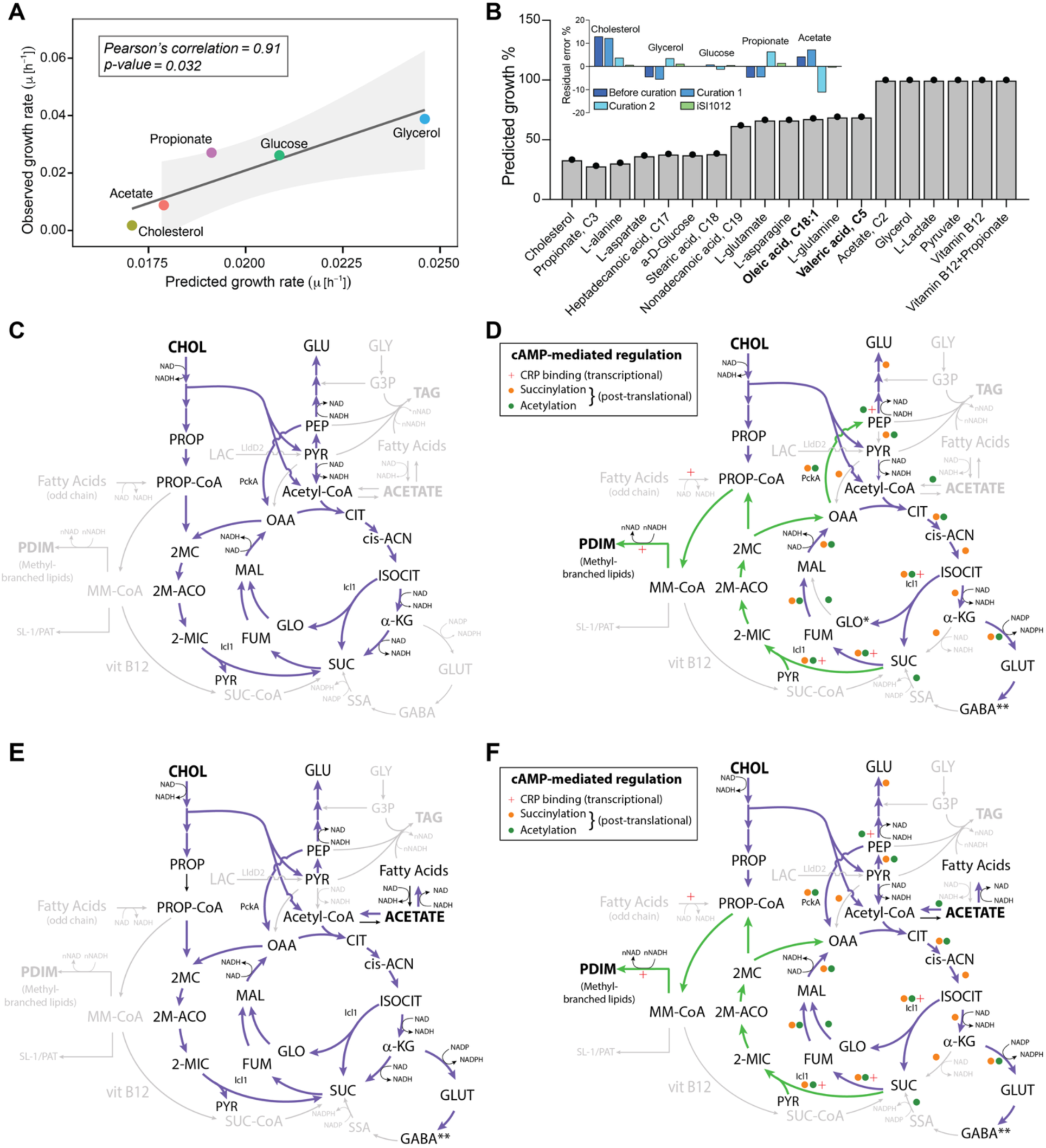
Contextualized iSI1012 model-predicted phenotypes and flux states of Mtb with and without drug treatment in inhibitory and non-inhibitory conditions. A. Correlation between model-predicted versus observed growth rate using media-constrained iSI1012 metabolic model. B. Model-predicted growth rescue by supplementing additional carbon sources in cholesterol-drug treated GIMME model. Inset shows residual prediction errors before and after multiple iterations of metabolic model curation in experimentally-tested contexts. Flux states of CCM during Mtb growth with and without drug treatment in inhibitory (C and D) and non-inhibitory (E and F) conditions. Gray arrows and associated metabolites indicate reactions with minimal to no flux; **purple arrows** indicate similar flux with and without drug treatment, whereas **green arrows** indicate treatment-induced reversal of directionality of flux. Inset key indicates color code and symbols for reactions catalyzed by enzymes that are known targets of cAMP-mediated transcriptional red plus (**+**) and post-translational succinylation orange dots (.) and acetylation green dots (.) based regulation.

**Table 1.** Curation of metabolic model.

| Details | iEK1011<br>model<sup>71</sup> | Curated<br>version 1 | Curated<br>version 2 | iSI1012 | Reference |
| --- | --- | --- | --- | --- | --- |
| **Flux predictions** |  |  |  |  |  |
| Reversal of MCC<br>(Lactate as carbon source) | - | x | x | x | Serafini et al 2019<sup>36</sup> |
| Reversal of MCC<br>(Pyruvate as carbon source) | - | x | x | x | Serafini et al 2019<sup>36</sup> |
| Reversal of MCC<br>(Glycerol as carbon source) | - | x | x | x | Borah et al 2021<sup>72</sup> |
| Forward MCC<br>(Cholesterol as carbon source) | x | x | x | x | Borah et al 2022<sup>72</sup> |
| Forward MCC<br>(Propionate as carbon source) | x | x | x | x | Koh and Rhee 2014<sup>29</sup> |
| Oxidative TCA cycle<br>(not FRD as major contributor) | - | - | x | x | Chang and Guan<br>2021<sup>2</sup> |
| CytBD is used when Cyt bcc is<br>compromised | - | - | - | x | Kalia et al 2019<sup>18</sup> |
| Type 2 Ndh is used when type I<br>(Nuo) is inhibited | - | - | - | x | Beites et al 2019<sup>44</sup> |
| **Reaction curation** |  |  |  |  |  |
| Removal of LDH reaction with<br>NADH as cofactor | - | - | x | x | Billig et al 2017 and<br>Stanley et al 2024<sup>37,73</sup> |
| Removal of Rv0694 from the LDH<br>reaction (i.e., LldD1 removal) | - | x | x | x | Billig et al 2017 and<br>Stanley et al 2024<sup>37,73</sup> |
| Removal of Rv1127c from PPDK<br>reaction | - | x | x | x | Borah et al 2021<sup>72,74</sup> |
| Electron and proton assignments | - | - | x | x | McNeil et al 2022 and<br>Cook and Poole<br>2016<sup>40,76</sup> |
| CytBD is non-proton pumping | - | - | x | x |  |
| Couple succinate oxidation to the<br>reduction of menaquinone | - | - | - | x | Adolph et al 2022<sup>123</sup> |
| Couple FRD to the reduction of<br>menaquinone | - | - | - | x | Adolph et al 2022 and<br>Watanabe et al<br>2011<sup>31,123</sup> |
| GPR for FRD and SUCCDi | - | - | x | x | Adolph et al 2022 and<br>Safarian et al<br>2021<sup>75,123</sup> |
| GPR of CytBD | - | - | x | x |  |
| FPRA reaction mass balanced | - | - | x | x | Model stoichiometry |
| Acetate transport (Act2r) as symport | - | - | x | x | Model stoichiometry |
| Reactions ETF and ETFD to re-<br>oxidize FADH2 to ubiquinone | - | - | x | x | Beites et al 2021<sup>44</sup> |
| **Growth predictions** |  |  |  |  |  |
| Growth on Cholesterol | x | x | x | x | Pandey et al 2008<br>and<br>Griffin et al 2012<sup>7,25</sup> |
| Growth on Glycerol | x | x | x | x | Pandey et al 2008<br>and<br>Griffin et al 2012<sup>7,25</sup> |
| Growth on Glucose | x | x | x | x | Billig et al 2017<sup>73</sup> |
| Growth on Acetate | x | x | x | x | Rucker et al 2015<sup>124</sup> |
| Growth on Propionate | x | x | x | x | Mulholland et al<br>2024<sup>91</sup> |
| Growth on Lactate via reverse MCC | - | x | x | x | Billig et al 2017 and<br>Stanley et al 2024<sup>37,73</sup> |
| Growth on Pyruvate via reverse<br>MCC | - | x | x | x | Serafini et al 2019<sup>36</sup> |
| Total no. of reactions curated | - | 54 | 56 | 62<br>(8 GPR) | Curation details |
| x = Curated/updated; - = Not accurate; GPR = Gene Protein Reaction Relationship |  |  |  |  |  |

Interestingly, across both inhibitory and non-inhibitory conditions, the “drug-treated” models derived using transcriptomes by GIMME^83^ algorithm revealed that GSK286 treatment resulted in strikingly similar remodeling of CCM, with a distinct hallmark reversal of MCC (**Figure 2 C-F)**. This finding revealed that drug treatment results in a blockade in the complete utilization of cholesterol, with propionyl Co-A, one of the principal breakdown products of cholesterol, redirected away from biomass production via CCM, and towards PDIM biosynthesis (**Figure 2D and 2F, highlighted in green)**. However, the model predicted that acetyl Co-A and pyruvate, intermediates produced in earlier steps of cholesterol breakdown, continued to support partial biomass production through supply of carbon into CCM, explaining why GSK286 treatment does not completely inhibit Mtb growth on cholesterol.

While the drug treatment resulted in similar remodeling of CCM in non-inhibitory conditions, there were some notable differences, including the maintenance of flux towards the TCA cycle through the reaction catalyzed by pyruvate carboxykinase (PckA) (**Figure 2F)**. By contrast, drug treatment in inhibitory conditions resulted in reversal of this reaction toward gluconeogenesis (**Figure 2D)**. Thus, the model predicted that supplementation with acetate restored carbon flux into CCM, relieving growth inhibition caused by the drug-induced blockade of cholesterol utilization. A noteworthy finding was that the context-dependent reversal of flux through reactions catalyzed by gene products of *icl1* and *pckA* was not obvious at the transcriptional level, considering that both genes were significantly upregulated across both inhibitory and non-inhibitory conditions (**Figure 1F and 1G**). Also notable was the finding that both Icl1 and PckA are known targets of post-translational regulation by cAMP-mediated acetylation and succinylation^86,87^. A systematic assessment uncovered that the 473 reactions through which flux was significantly altered (i.e., increased, decreased, reversed, or blocked; paired t-test p-value < 0.05) by GSK286 treatment were catalyzed by enzymes enriched for targets of cAMP-mediated post-translational regulation by acetylation^86^ (*p-value = 4.6e-05*) and succinylation^87^ (*p-value = 0.0033*; **Data S2**). In sum, analysis of flux states and FBA using the metabolic model identified a blockade in cholesterol breakdown due to the reversal of the MCC pathway as a plausible mechanistic explanation for the conditional growth inhibition of Mtb by Rv1625c agonists.

### Restoration of carbon flux with alternate growth substrates reduces the efficacy of Rv1625c agonists

Interestingly, simulations with both transcriptome-contextualized and the base iSI1012 models predicted that, similar to acetate, other substrates that restore carbon flux into CCM, specifically the TCA cycle, through alternate routes would support continued biomass production and thereby reduce growth inhibitory consequences of Rv1625c agonists (**Table S1**). Consistent with model predictions, growth inhibition of Mtb by GSK286 (**Figure 3A**) and TBD11 (**Figure 3B**), was reversed entirely or partially in glycerol, acetate, lactate, pyruvate, oleic acid, and other substrates, provided individually or in combination with cholesterol (**Figure 3, Figure S1, and Table S2**). The two outliers, i.e., partial rescue by oleic acid supplementation (both drugs) and continued growth inhibition by TBD11 in cholesterol + pyruvate, are addressed in later sections.

**Figure 3.**
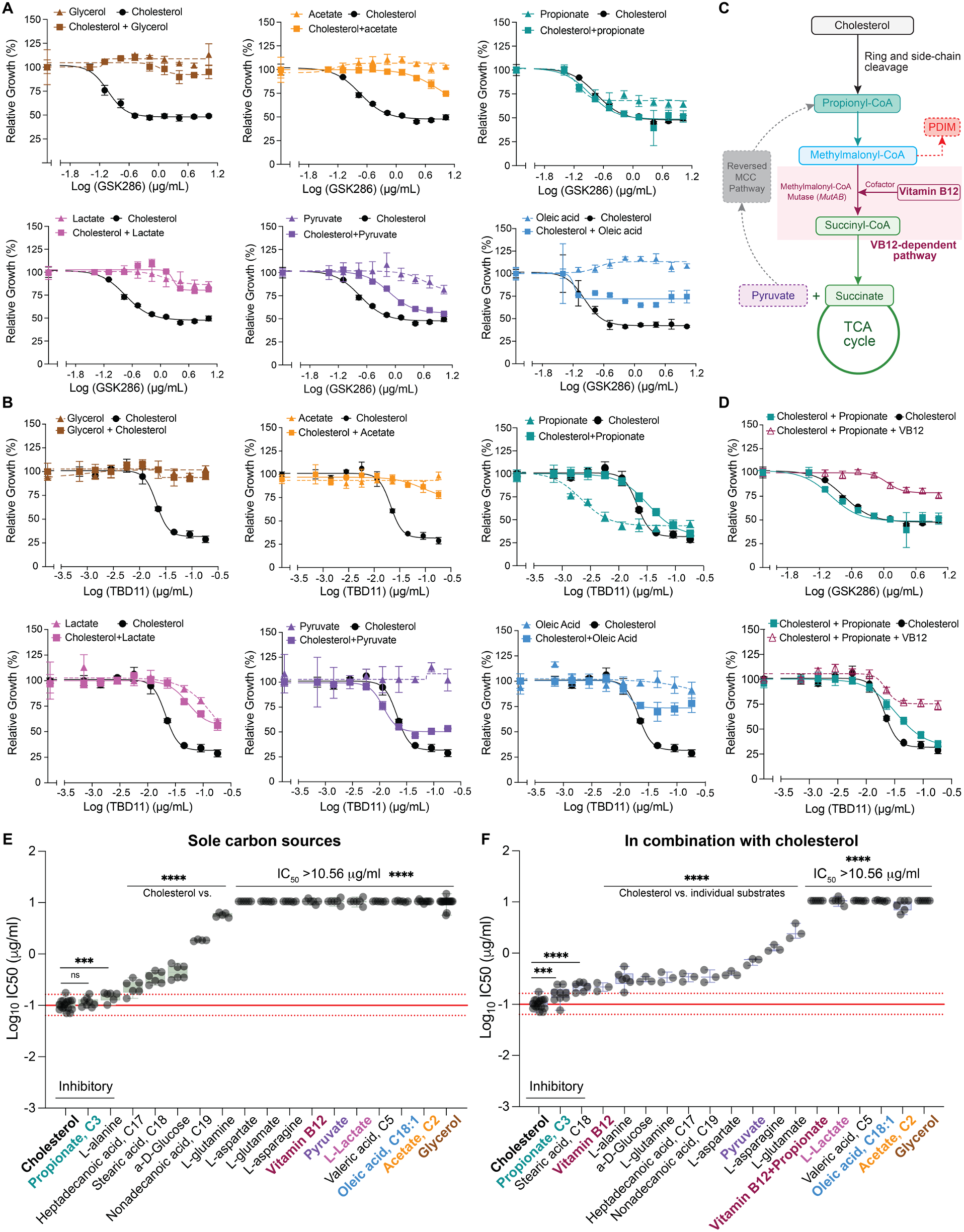
Supplementation with nutrients that restore carbon flux into CCM reduce sensitivity to Rv1625c agonists. Mtb growth inhibition by GSK286 (**A**) and TBD11 (**B**) in Sauton’s minimal media supplemented with glycerol, acetate, lactate, pyruvate, oleic acid, and propionate. **C**. Pathway schematic of metabolic flux state of Mtb with vitamin B12 supplementation to replenish TCA cycle by activating consumption of propionyl Co-A via the methyl malonyl-coA pathway. **D**. Relative growth inhibition by GSK286 and TBD11 upon addition of vitamin B12 (VB12). Distribution of IC50 of GSK286 in the presence of each substrate as a sole carbon source (**E**) and in combination with cholesterol (**F**). P-values are deduced using two-way ANOVA, where **<0.01, ***<0.001, and ****<0.0001. The dose-response curve in cholesterol shown in each plot is from the corresponding experiment batch. Since many conditions were tested in the same batch, the dose-response curve for cholesterol was duplicated across figure panels from the same experiment. Data are representative of at least two independent experiments. Error bars correspond to standard deviation of replicates within the same experiment.

In contrast, propionate, a cholesterol breakdown product, did not alleviate Mtb growth inhibition by Rv1625c agonists (**Figure 3**), individually or with cholesterol, further supporting the hypothesis that growth inhibition by these drugs is likely due to the reversal of MCC, which blocks carbon flux towards biomass production. Additional support for this hypothesis came from the observation that Rv1625c-induced growth inhibition in cholesterol + propionate was also abolished upon addition of vitamin B12 (VB12), a cofactor of methylmalonyl CoA mutase (*MutAB*), which enables conversion of propionate into succinate via the methylmalonyl CoA pathway, thereby restoring carbon flux into the TCA cycle (**Figure 3C and D**). In sum, systematic survey of drug efficacy across diverse nutritional contexts, confirmed model-predicted conditional activity of Rv1625c agonists with a log-fold increase in IC50 in non-inhibitory conditions relative to inhibitory conditions (**Figure 3E-F, Figure S1, Table S2**).

### Trade-off between PDIM biosynthesis and biomass production underlies condition-dependent drug activity

In addition to uncovering how alternate substrates can counter MCC reversal and restore carbon flux into CCM, the model identified that the context-dependent efficacy of Rv1625c agonism is governed by a fundamental metabolic trade-off between virulence lipid synthesis and biomass production. FBA, using robustness analysis^88^, revealed competition for key carbon precursors between these two demands across both inhibitory and non-inhibitory conditions. Specifically, PDIM biosynthesis, which is essential for Mtb virulence and cell wall integrity, requires substantial quantities of propionyl-CoA, methylmalonyl-CoA, and NAD(P)H, precursors that also serve as inputs to biomass-generating pathways. A strong negative correlation between predicted biomass yield and PDIM flux confirmed that increase in PDIM production comes at a direct cost to growth (**Figure 4A**).

**Figure 4:**
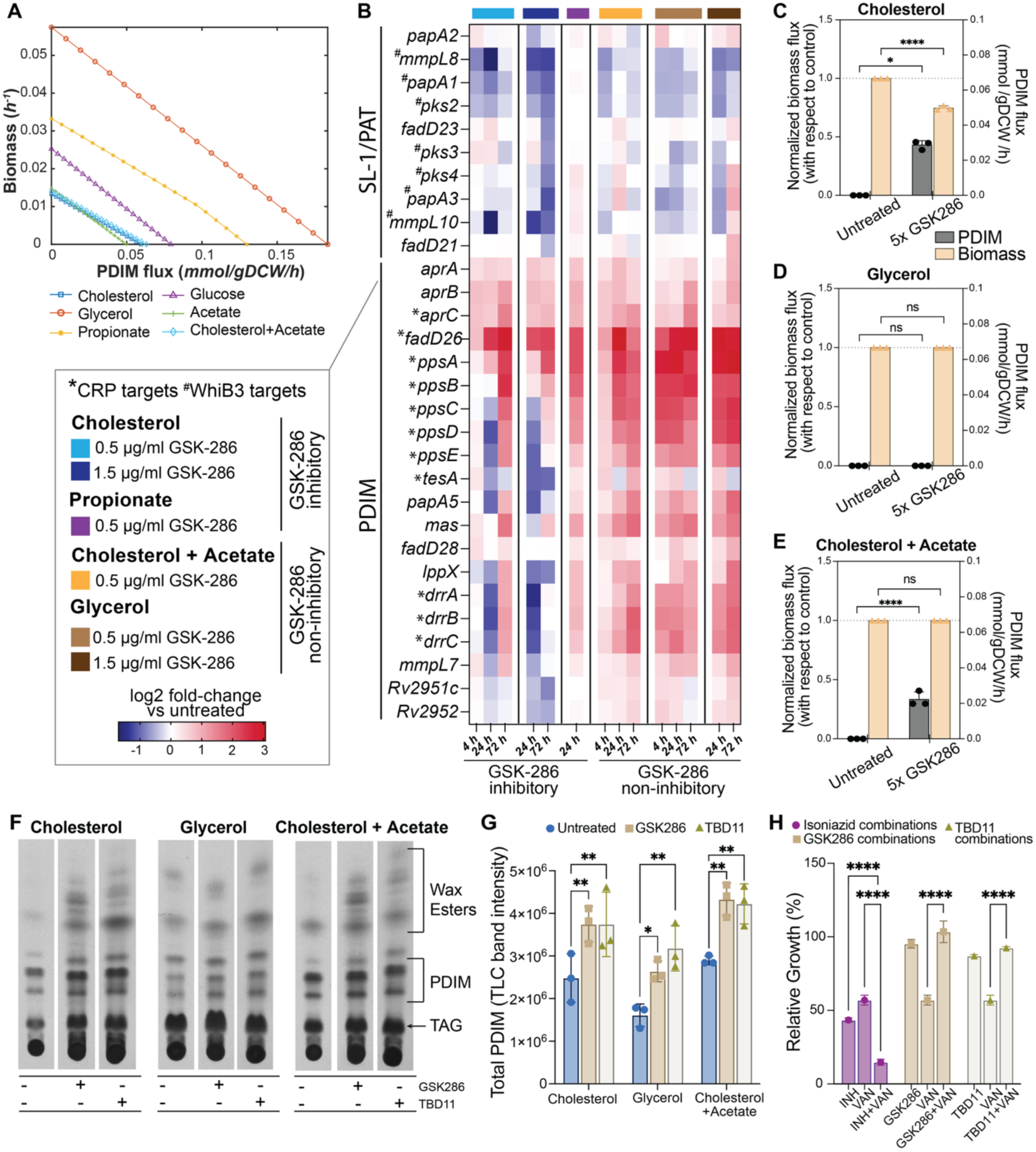
Role of PDIM biosynthesis in condition-dependent growth inhibition. **(A)** *In silico* model simulations predict trade-off between biomass production and flux towards PDIM biosynthesis. **(B)** Heatmap illustrates differential expression of PDIM biosynthetic genes across treatment conditions. CRP-regulated targets are indicated with an asterisk (*). Model predicted biomass and PDIM levels with biomass production as the objective function for FBA in cholesterol (**C**), glycerol (**D**) and cholesterol + acetate (**E**). (**G**) TLC radiograms and corresponding densitometry bar plots (**H**) for PDIM levels in Mtb cultures in cholesterol, glycerol, and cholesterol + acetate at 72hrs with and without treatment with GSK286 and TBD11. The TLC plates were loaded with equivalent biomass of Mtb (normalized by wet cell weight) across all conditions. (**H**) Relative growth inhibition of Mtb cultures in 7H9+GOT treated with vancomycin alone and in combination with Rv1625c agonists or isoniazid. Data are representative of at least two independent experiments. Error bars correspond to standard deviation of replicates within the same experiment. P-values are deduced using two-way ANOVA, where **<0.01, ***<0.001, and ****<0.0001.

To test this model prediction at the transcriptional level, we examined the expression of PDIM biosynthetic pathway genes under drug treatment. Strikingly, the entire PDIM gene cluster was upregulated in response to GSK286 across both inhibitory and non-inhibitory conditions. By contrast, GSK286 treatment resulted in the downregulation of sulfolipid (SL-1) and polyacyl trehalose (PAT) biosynthetic pathway genes, including direct regulatory targets of WhiB3^89^, across both inhibitory and non-inhibitory conditions (**Figure 4B**). This finding demonstrates that transcriptional drive toward virulence lipid synthesis is a primary, condition-independent consequence of Rv1625c agonism. Consistent with the broader transcriptional remodeling described above, 11 out of 20 differentially expressed PDIM genes are regulated by CRP (*p-value = 4.78e^-^*^10^), establishing a direct regulatory link between drug-induced cAMP elevation and PDIM pathway activation. Interestingly, 9 out of the 20 enzymes are also targets of cAMP-mediated post-translational regulation by acetylation^86^ or succinylation^87^ (hypergeometric enrichment *p-value = 1.49e^-0^*^7^, **Data S2**).

To explore consequences of this global transcriptional remodeling on growth and PDIM production, we performed FBA using biomass production as the primary objective function across inhibitory (**Figure 4C**) and non-inhibitory conditions (**Figure 4D and E**). In inhibitory, cholesterol-only condition, forcing the metabolic network to optimize for biomass production in the presence of drug, still resulted in a diversion of carbon flux toward PDIM pathways (**Figure 4C**). This diversion of carbon flux prevents the network from achieving normal growth yields under drug treatment in cholesterol. Importantly, the residual biomass production simulated on cholesterol recapitulated experimental observations of incomplete growth inhibition in cholesterol (**Figure 3**). This residual growth reflects the ability of metabolic intermediates, specifically pyruvate and acetyl-CoA that are produced in early steps of cholesterol breakdown, to enter CCM via alternate routes and partially bypass the primary metabolic blockade by reversal of MCC and supply intermediates to essential downstream pathways. Conversely, under non-inhibitory conditions, particularly when cholesterol is supplemented with acetate, the blockade in biomass production is largely lifted (**Figure 4E**). Acetate supplementation replenishes the central metabolic network with two-carbon units, satisfying the energetic demands of both PDIM synthesis and essential biomass pathways simultaneously, thereby restoring normal growth. In contrast to the protective effect of acetate, our model predicted that propionate supplementation would exacerbate the toxicity of the drug by loading the cell with three-carbon precursors that feed directly into the virulence lipid synthesis branch points. This prediction was borne out by the enhanced growth inhibition observed with TBD11 under propionate conditions (**Figure 3B**). Collectively, the model predictions account for the conditional activity of Rv1625c agonists, rationalizing the dramatic, log-fold difference in IC50 of GSK286 between the inhibitory and the non-inhibitory conditions (**Figure 3E-F, Table S2**). Thin-layer chromatography (TLC) analysis directly confirmed these metabolic model predictions at the phenotypic level, showing significant accumulation of PDIM and related surface lipids (e.g., wax esters^90^) in treated cells from both inhibitory and non-inhibitory conditions (**Figure 4F and G**). Consistent with the drug-induced increase in PDIM, which is known to reduce cell-wall permeability for higher molecular weight antibiotics^91^, Rv1625c agonist treatment rendered Mtb more resistant to vancomycin: DiaMOND analysis of the agonist-vancomycin combination revealed strong antagonism (**Figure 4H, FIC>2**). Together, these findings provide orthogonal, phenotypic confirmation that drug treatment remodels the cell wall through elevated PDIM production.

### Genetic perturbations of key metabolic branch points validate model-predicted mechanism of condition-dependent drug activity

Our model predicted that drug sensitivity is dictated not merely by the magnitude of carbon flux, but by the specific rerouting of that flux through CCM. To validate this, we utilized genetic perturbations to manipulate key metabolic branch points, testing whether altering carbon flux paths could predictably modulate condition-dependent drug activity. We investigated key reactions catalyzed by Icl1 and PckA, which gate flux between the TCA cycle, MCC, the anaplerotic route, gluconeogenesis, and lipid synthesis. First, CRISPRi knockdown (KD) recapitulated essentiality of *icl1* for growth of Mtb on cholesterol ^25,27,29,30^ (but not glycerol), and the essentiality of *pckA* in both growth contexts^68–70^ (**Figure 5A and B**). Second, both *icl1* KD and *pckA* KD effectively abolished drug activity in cholesterol (**Figure 5A**). Intriguingly, in glycerol-supplemented media, *icl1* KD resulted in significant growth improvement during drug treatment (**Figure 5A lower panel**). The *icl1* KD phenotype was consistent with the model prediction that blocking the glyoxylate shunt reduces the misdirected carbon flux toward PDIM synthesis, and instead reroutes glycerol toward biomass synthesis. Collectively, these findings of Icl1 and PckA (**Figure 5 A and B**) validate the metabolic network model, demonstrating that the reversal of Icl1- and PckA-catalyzed reactions at these branch points are key components of the MoA of Rv1625c agonists.

**Figure 5:**
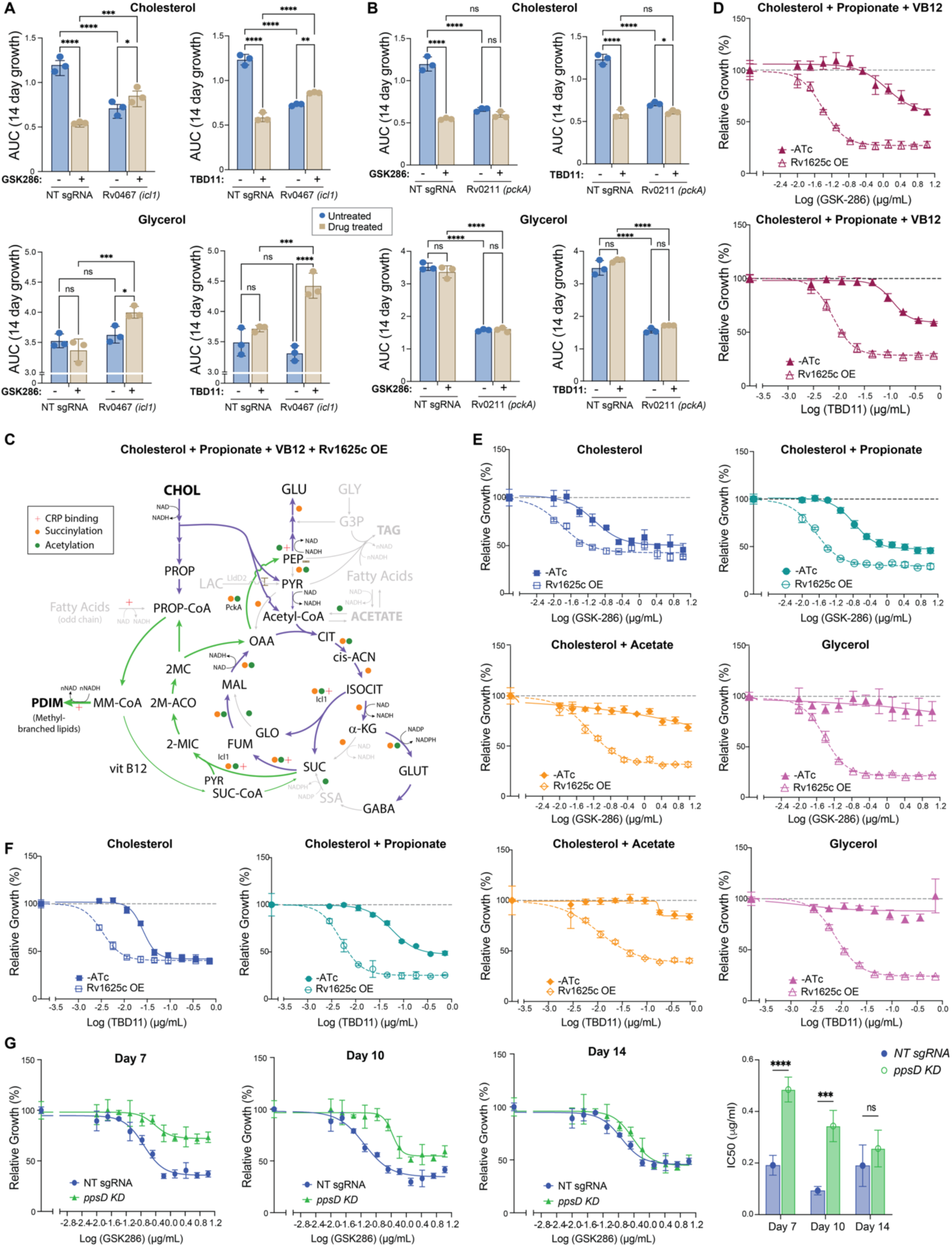
Genetic perturbations of key metabolic branch points modulate condition-dependent drug efficacy. Condition-dependent effects of CRISPRi KD of *icl1* (A) and *pckA* (B) on relative growth inhibition of Mtb by TBD11 and GSK286. Bar plots indicate cumulative biomass (AUC) of WT and KD strains in cultures without (-) or with (+) with GSK286 or TBD11 treatment. P-values were deduced using two-way ANOVA, where **<0.01, ***<0.001, and ****<0.0001. **C**. Pathway schematic shows predicted changes upon Rv1625c OE in cholesterol + propionate conditions supplemented with VB12. **D**. Relative growth inhibition of Mtb by GSK286 (top) and TBD11 (bottom) in VB12 supplemented media with (+ATc) and without (-ATc) induction of Rv1625c OE. Relative growth inhibition of Mtb by GSK286 (**E**) and TBD11 (**F**) across inhibitory and non-inhibitory carbon sources with (+ATc) and without (-ATc) induction of Rv1625c OE. Data are representative of at least two independent experiments. Error bars correspond to standard deviation of replicates within the same experiment. (G) Relative growth inhibition of Mtb in dose-response assays with GSK286 in MSM with cholesterol, with and without CRISPRi KD of *ppsD*. The bar plot shows IC50 of GSK286, derived from the dose-response assays.

Consistent with our finding that drug treatment causes similar global metabolic remodeling across all conditions (**Figure 2**), Rv1625c overexpression (OE) potentiated GSK286 and TBD11 activity in inhibitory conditions^58,59^ and restored their activity even in non-inhibitory conditions, including in propionate + VB12 (**Figure 5C-F**). This finding supports the model’s assertion that the mechanism of action of Rv1625c is not cholesterol-specific and a consequence of diverting carbon flux away from biomass production and toward PDIM synthesis, which can be accomplished by Rv1625c OE across all growth contexts. Indeed, CRISPRi knockdown of *ppsD*, a CRP-regulated polyketide synthase essential for PDIM biosynthesis^92^, significantly reduced growth inhibition by Rv1625c agonists over 10 days of drug treatment, indicating that flux into PDIM is required for the growth-inhibitory effect (**Figure 5G**, see Discussion).

### Rv1625c agonism involves ETC impairment and redox imbalance

Having established that Rv1625c agonists redirect carbon flux toward PDIM biosynthesis and away from biomass production, we next investigated the upstream metabolic trigger responsible for this remodeling. Transcriptome analysis revealed that among upregulated genes in non-inhibitory conditions, there was a significant enrichment of genes encoding NAD(P)H-utilizing enzymes (*p-value = 0.00032*; **Data S3**), suggesting a systemic response to disrupted cellular redox balance. This observation raised two non-exclusive hypotheses: first, that dissipation of reductive stress may underlie the absence of growth inhibition in non-inhibitory conditions; and second, that carbon flux entering the TCA cycle through alternate routes supports continued growth and respiration, buffering the redox consequences of drug action.

Consistent with these hypotheses, drug treatment was associated with a coordinated shift in expression of genes associated with the respiratory machinery across all conditions (**Figure 6A**). Most notably, genes of the *nuo* operon which encode the respiratory type I NADH dehydrogenase (NDH-1) complex were significantly downregulated in both inhibitory and non-inhibitory conditions (**Figure 6A**), while *ndh* (*Rv1854c*), a CRP-regulated gene encoding the type-II NADH dehydrogenase NDH-2, was markedly upregulated across all conditions tested but not *ndhA (Rv0392c)* (**Figure 6B**). This reciprocal regulation is consistent with prior reports that NDH-1 blockade induces NDH-2 as a compensatory mechanism^44–46^. Transcriptome-contextualized flux models corroborated these transcriptional findings by uncovering that drug treatment resulted in near-complete blockade of flux through NDH-1 and a compensatory increase in flux through NDH-2 across both inhibitory and non-inhibitory conditions. Additionally, drug treatment in cholesterol conditions shifted cytochrome usage from the bc₁-aa₃ branch toward cytochrome bd, reflecting a broader reorganization of respiratory electron flow under drug pressure (**Figures 6C** and **6D, Data S2**).

**Figure 6:**
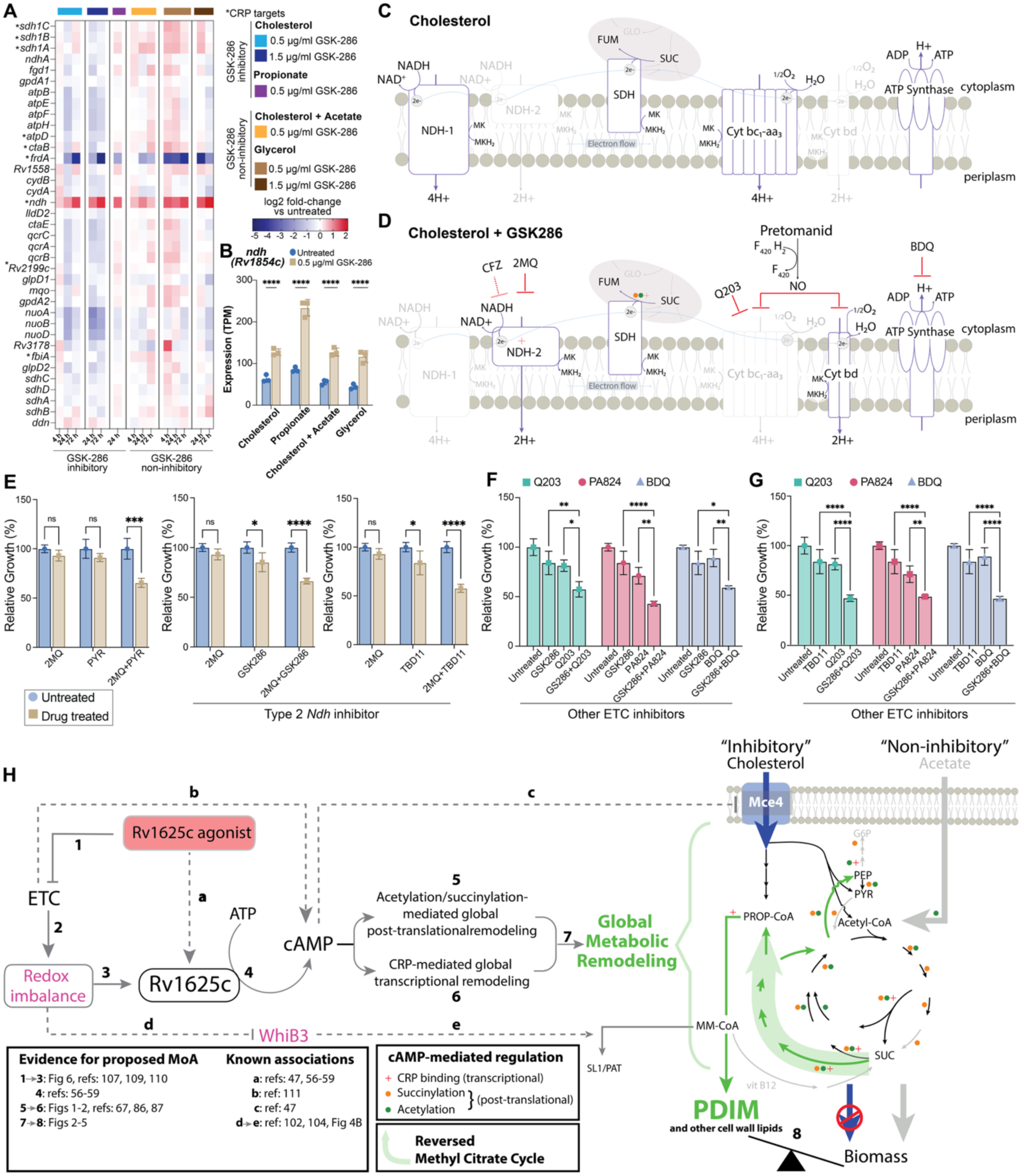
ETC impairment by 1625c agonists and their associated synergistic drug combinations. **A**. Heatmap shows log-fold expression level changes of genes encoding components of respiration and ETC in response to 5X and 15X GSK286 treatment across inhibitory (cholesterol and propionate) and non-inhibitory conditions (cholesterol + acetate and glycerol) (see inset key for details). **B**. Bar plots show *ndh* (Rv1854c) expression levels without and with GSK286 treatment across the same conditions as in (A). FBA-predicted flux states of the electron transport chain under baseline (**C**) and drug-treated conditions (**D**) in cholesterol. Dark outlines of ETC components and dark arrows indicate active pathways with significant flux values, whereas gray outlines and arrows indicate pathway components with negligible flux. ETC targeting drugs and their targets are shown in GSK286-treated ETC flux map in panel D. **E-G**. Bar plots illustrating relative growth inhibition of Mtb in MSM with cholesterol by individual and two-way drug combinations of GSK286 and TBD11 with 2MQ and other ETC inhibitors. **H.** Schematic for likely mechanism of action of Rv1625c agonists. The agonists cause a blockade in the ETC resulting in redox imbalance, which is the likely trigger that activates Rv1625c. The elevated cAMP levels drive global remodeling of metabolism at the transcriptional level via CRP, and post-translationally through acetylation/succinylation of key CCM enzymes. The global metabolic remodeling results in the reversal of MCC, which redirects the breakdown product of cholesterol (propionyl CoA) away from biomass production. A simultaneous transcriptional and post-translational reduction in WhiB3 reduces flux towards SL1/PAT, diverting methylmalonyl CoA towards cell wall lipids, including PDIM (evidence supporting each claim presented in this study or in literature are indicated; dashed lines indicate previously reported associations that are consistent with the model).

To integrate these findings quantitatively, we used contextualized models where transcriptome data were available, and base models with appropriate media constraints otherwise, to predict intracellular NADH levels across all inhibitory and non-inhibitory conditions. Model-predicted NADH levels were significantly correlated with experimentally determined IC50 values (*Pearson p-value = 0.0002267, Spearman p-value = 0.0003246*; **Figure S2**), supporting the interpretation that reductive stress is a key determinant of drug efficacy. A log-normal fit to the NADH versus IC50 relationship suggested threshold-dependent dynamics, i.e., that NADH likely activates Rv1625c when it accumulates beyond a critical threshold^93^. Three substrates emerged as outliers from this correlation: propionate, alanine and oleic acid. Propionate feeds directly into the reversed MCC, explaining why the drug is effective with this substrate even though the reductive stress is low. Alanine metabolism bypasses early glycolytic and TCA cycle control points through direct transamination to pyruvate, decoupling its NADH contribution from that predicted by CCM^94^. Oleic acid, which enters metabolism via β-oxidation and generates high levels of FADH₂ and acetyl-CoA, similarly perturbs the expected NADH/NAD⁺ ratio^95^. However, oleic acid has also been found independently to directly inhibit Rv1625c^61^, providing an additional explanation for its partial rescue phenotype.

These observations prompted us to evaluate whether the drug-induced blockade in the ETC could be exploited for synergistic combination therapy. The drug-treated flux maps suggested synergistic interactions of Rv1625 agonists with other established ETC inhibitors (**Figure 6D**). DiaMOND assays^96,97^ under fixed drug ratios (**Figure 6E-G, S3, and S4**), validated model predictions by demonstrating synergy of Rv1625c agonists with bedaquiline (ATP synthase inhibitor)^98^, pretomanid (inhibits both cytochromes)^99^, and Q203 (cytochrome bcc:aa_3_ inhibitor)^100^. The most striking finding was the synergy of Rv1625c agonists with 2-mercapto-quinazolinone (2-MQ; DDD00853663)^42^, a specific inhibitor of the NDH-2 complex, which was consistent with co-engagement of the NDH-2-dependent compensatory pathway activated by drug treatment (**Figure 6E**). Furthermore, partial overlap in both activity and synergy profiles with NDH-I inhibitors pyridaben and rotenone reinforced the centrality of NDH-1 blockade to the mechanism of action of Rv1625c agonists. However, the Rv1625c-independence of growth inhibition by pyridaben and rotenone (**Figure S3**) suggested that Ndh-1 blockade by GSK286 and TBD11 occurs via a distinct and novel mechanism of action.

## Discussion

The dependence of drug activity on host-relevant carbon sources presents a formidable challenge to conventional drug discovery, yet it offers a novel perspective into the coupling of bacterial physiology, cellular energetics, and transcriptional networks. In this study, by integrating temporal transcriptomics with an updated genome-scale metabolic model (iSI1012) and targeted genetic perturbation, we elucidated the metabolic underpinnings of the context-dependent activity of Rv1625c agonists, GSK286 and TBD11. Rather than acting through classical disruption of an essential housekeeping enzyme, these agonists effect global metabolic reprogramming by hijacking an endogenous second-messenger switch and cascade to reroute carbon away from biomass production and towards virulence-lipid production.

A central finding of our study is that the global, cAMP-driven transcriptional program elicited by Rv1625c activation remains remarkably uniform, irrespective of whether the external carbon substrate permits (inhibitory) or prevents (non-inhibitory) growth inhibition by the adenylate cyclase agonists. Exposure to GSK286 resulted in transcriptome-wide changes characterized by a distinct regulatory signature of the cAMP receptor protein (CRP). Core anaplerotic and central metabolic nodes, including *icl1* and *pckA*, were significantly upregulated under both inhibitory (cholesterol, propionate) and non-inhibitory (cholesterol + acetate, glycerol) carbon conditions. This decoupling of the global transcriptional response from the downstream phenotypic outcome indicates that the nutrient-dependent activity of Rv1625c agonists is not driven by differential gene expression. Instead, it is governed by the flux capabilities of the available nutrient pools. Furthermore, our finding that the enzymes catalyzing these carbon flows are significantly enriched for known post-translational modifications (PTMs), specifically cAMP-mediated acetylation and succinylation, suggests a multi-layered regulatory architecture that simultaneously drives CRP-dependent transcription while altering flux partitioning.

By updating and contextualizing the constraints-based metabolic network model iSI1012, we discovered that energetic and carbon-precursor trade-off between PDIM biosynthesis and biomass production is a centerpiece of the conditional toxicity of Rv1625c agonists. The activation of the PDIM biosynthetic cluster acts as a metabolic sink, draining the intracellular pool of vital multi-carbon building blocks (acetyl-CoA, propionyl-CoA, and malonyl-CoA) alongside substantial reducing equivalents (NADH). When Mtb is restricted to cholesterol or propionate as its primary carbon source, the breakdown pathways channel carbon directly through the MCC. Drug action in these conditions forces a reversal of the MCC, driving cholesterol-derived propionyl-CoA directly to PDIM synthesis. This metabolic diversion deprives the cell of the precursors needed to manufacture essential biomass components and amino acids, stalling biomass production as well as reducing cholesterol uptake (**Figure 6H**). Importantly, our flux simulations successfully recapitulate the partial growth inhibition observed experimentally on cholesterol, attributing it to early steps in cholesterol side-chain cleavage that produce acetyl-CoA and pyruvate, which partially bypass the primary metabolic blockade and replenish baseline CCM to preserve a low-level, residual rate of biomass synthesis. Conversely, this biomass burden is fully mitigated under non-inhibitory conditions, such as acetate or glycerol supplementation, in which the alternative substrates replenish the TCA cycle with two- and three-carbon units. This influx satisfies the intensive precursor and energetic demands of PDIM synthesis while maintaining the carbon flux required to sustain gluconeogenesis and the TCA cycle for biomass production. This model was further validated by genetic perturbation: single gene knockdowns of *icl1* and *pckA* block these critical metabolic branch points, cutting off the downstream flow of misdirected carbon and alleviating drug activity in cholesterol as well as glycerol. Most notably, the finding that *icl1* knockdown in glycerol-supplemented media actually improves bacterial growth under drug treatment provides strong evidence for our model, in which blockade of the glyoxylate shunt spares glycerol from diversion to lipid production and redirects it toward biomass.

It was noteworthy that, as predicted by our model, *ppsD* knockdown reduced drug activity, but surprisingly this effect was confined to the first ∼10 days of treatment. Our TLC analysis offers a plausible explanation by revealing that drug treatment also resulted in the accumulation of additional (yet to be characterized) cell-wall lipids whose biosynthesis may likely draw on the same methylmalonyl-CoA pool^101^; over time, methylmalonyl-CoA blocked from PDIM synthesis by *ppsD* knockdown may be rerouted into these other lipids, restoring growth inhibition and explaining why the reduction in drug activity is only transient. Alternatively, or in parallel, blocking PDIM synthesis removes a major sink for cholesterol-derived carbon, which would be expected to cause a buildup of upstream breakdown products and PDIM precursors, including propionyl-CoA, a metabolite known to be toxic to Mtb^91,102^. In this case, the eventual restoration of growth inhibition may occur through a mechanism distinct from that in PDIM-competent cells - driven by the toxicity of accumulated intermediates rather than by diversion of carbon flux away from biomass production.

A striking feature of this trade-off is that the activation of PDIM synthesis may be not merely a consequence of carbon rerouting but an adaptive response to the redox state of the cell^89,91,103,104^. Because the synthesis of methyl-branched virulence lipids consumes large quantities of reducing equivalents, the drug-induced upregulation of the PDIM cluster may represent an attempt by Mtb to dissipate reductive stress through lipid anabolism^89,91,103,104^. This interpretation directly links the PDIM trade-off to our redox findings (below): the same NADH accumulation that we propose activates Rv1625c could also be partially relieved by channeling carbon and reducing power into PDIM, making lipid anabolism both a symptom of, and a buffer against, the underlying redox imbalance.

At the upstream end of the cAMP cascade, our data indicate that Rv1625c activation is localized to the ETC, driven potentially by an apparent functional blockade of the NDH-1 complex. Drug exposure causes transcriptional downregulation of the *nuo* operon, accompanied by strong upregulation of *ndh* as well as genes encoding alternative NAD(P)H-utilizing enzymes, a signature of a global effort to rebalance the internal redox pool. The significant correlation between model-predicted intracellular NADH accumulation and experimental IC50 values across diverse carbon sources (**Figure S2**) strongly supports a second-messenger role for cellular redox status, in which the NADH/NAD⁺ ratio, and thus the state of the ETC, gates Rv1625c expression and activation. This framework also offers a parsimonious explanation for why the drugs are inactive in non-inhibitory conditions, where Rv1625c induction may simply not reach a sufficient threshold. Two non-exclusive mechanisms could limit induction: first, substrates such as glycerol and acetate may not generate reductive stress to the degree that cholesterol does; and second, by restoring carbon flux into CCM through alternative routes, these substrates sustain biomass production and continued respiration, which further dissipates the reductive stress. Consistent with this threshold model, overexpression of Rv1625c is sufficient to override the protection afforded by these carbon sources: it forces larger amounts of carbon flux away from biomass and toward PDIM biosynthesis even in otherwise non-inhibitory conditions, restoring the inhibitory effect of drug treatment.

This state of respiratory electron blockade creates distinct vulnerabilities, which we confirmed using DiaMOND ^96,97^ synergy assays. The synergy observed between Rv1625c agonists and alternative ETC inhibitors, notably the NDH-2 inhibitor 2-MQ, corroborates that ETC impairment is central to Rv1625c agonism. Co-targeting distinct components of the ETC with compounds such as Q203, bedaquiline, and pretomanid is likewise synergistic (this study, as well as ^18,39,46,56,99,100,105^), accelerating redox imbalance while reducing ATP synthesis. Notably, the degree of synergy (**Figure S3**) between Rv1625c agonists and ETC inhibitors was inversely related to the molecular weight of the partner compound (**Figure S4**). We attribute this to a counteracting effect of the drug-induced cell-wall remodeling: while agonist treatment sensitizes Mtb to ETC inhibition, the concomitant increase in PDIM reduces cell-wall permeability to high-molecular-weight antibiotics, analogous to the PDIM-associated exclusion of vancomycin (**Figure 4H**), thereby limiting entry of larger inhibitors such as bedaquiline and Q203 (**Figure S3 and S4**). Pretomanid, a smaller compound whose uptake is unaffected by this remodeling, consequently shows the strongest synergy (**Figure 6F, G, S3, and S4**). In agreement with the recent study by Rabodoarivelo et al.,^106^ the triple drug combination of bedaquiline, pretomanid, and Rv1625c agonist exhibited robust synergy (**Figure S3**). However, in cholesterol-containing media, diagonal “fixed-ratio” measurements in our DiaMOND assays revealed that Rv1625c agonists interacted more synergistically with pretomanid than with bedaquiline, offering a complementary perspective to the view that bedaquiline is the primary driver. This apparent discrepancy between findings from the two studies may reflect differences in drug exposure and host niche. Rabodoarivelo et al.^106^ used substantially higher drug concentrations, which could mask the differential penetration we observe at lower doses. Moreover, bedaquiline is known to accumulate within host cells^107^; this intracellular enrichment may offset the reduced permeability imposed by Rv1625c-induced PDIM elevation, allowing bedaquiline to reach effective concentrations in the macrophage niche despite the cell-wall remodeling that limits its entry in axenic culture.

The synergy data also offer an insight into the precise mechanism underlying Rv1625c activation. Both pyridaben and rotenone synergize with NDH-2 inhibitor 2-MQ^46^, yet, unlike the Rv1625c agonists, their activity is independent of Rv1625c (**Figure S3**), suggesting that they impair NDH-1 through a mechanism distinct from that of the agonists. Pharmacological inhibition of NDH-1 (e.g., by pyridaben) is lethal to Mtb specifically through impaired NADH oxidation, since rescue by a water-forming NADH oxidase abolishes this toxicity^46^. This places the redox consequence, i.e., an inability to regenerate NAD⁺, at the center of NDH-1 inhibition, consistent with our model in which Rv1625c agonists act by generating reductive stress at NDH-1. One intriguing possibility raised by this contrast is that Rv1625c agonists impair NDH-1 specifically at the level of menaquinone reduction, either alone or in combination with an effect on NADH oxidation, thereby blocking electron flow at a step that uniquely generates the redox signal sensed by Rv1625c. The precedent we lean on is CyaC of *Sinorhizobium meliloti*^108^, a redox-regulated adenylyl cyclase with a quinone-responsive diheme-B membrane anchor, an example of a cyclase whose activity is gated by the redox/quinone state of the membrane, which is suggestive of how Rv1625c (also membrane-anchored^53,55,60^, likely heme-harboring^109^, and heme B inhibited^110^) might sense ETC status. Another study showed that the intracellular level of cAMP in *M. smegmatis* was found to be increased under ETC-inhibitory conditions^111^. Whether the agonists activate Rv1625c through direct binding or indirectly via redox state of the ETC, whether Ndh-1 blockade is upstream of cAMP or a downstream consequence of it, as well as resolving the exact biochemical target domain of the agonists within NDH-1 will be important directions for future work.

Finally, the conditional logic of these drugs may be deeply tied to the pathogenic lifestyle of Mtb itself. It offers a compelling explanation for why agonist treatment of non-pathogenic strains: *M. smegmatis*^57^ and *Rhodococcus jostii*^59^ engineered to express a heterologous copy of Rv1625c elevates cAMP yet fails to inhibit growth on cholesterol i.e., in the absence of the PDIM biosynthetic sink, the cAMP signal is uncoupled from the metabolic trade-off that drives toxicity. The Rv1625c-PDIM axis may therefore represent a circuit that evolved specifically in the context of host infection, linking the redox cues of the intracellular environment to the production of a key virulence lipid.

This tradeoff between growth inhibition and PDIM production, however, raises an important cautionary consideration that warrants careful evaluation before such agonists are advanced therapeutically. Because Rv1625c agonists actively drive flux into PDIM, a major virulence lipid, a critical open question is whether bacteria that survive sub-lethal or context-limited drug exposure are rendered more virulent, having been pharmacologically pushed toward a lipid-rich, immunomodulatory cell-envelope state. Understanding the immunological and pathogenic consequences of this forced virulence-lipid synthesis, particularly across the heterogeneous microenvironments Mtb occupies during infection, will be essential to determine whether the trade-off we describe is uniformly therapeutic or carries context-dependent risk.

In conclusion, this work shifts our understanding of conditional drug activity in Mtb from a reductionist view to a predictive, systems-level perspective that accounts for metabolic network topology. We have shown that Rv1625c agonists exploit an endogenous regulatory axis (**Figure 6H**), forcing a severe metabolic conflict between virulence-lipid synthesis and cell survival. From a translational perspective, these insights offer a clear roadmap for rational combination therapy: by pairing Rv1625c agonists with specific ETC inhibitors or metabolic adjuvants, it may be possible to leverage these global vulnerabilities to overcome the drugs’ conditional activity and arrest the pathogen in a destructive metabolic state regardless of the local microenvironment. Finally, our findings raise the possibility that the ∼15-17 adenylyl cyclases of Mtb act as a distributed network of sensors linking diverse cues to metabolic and cell-wall remodeling^47–53^, offering one explanation for the pathogen’s ability to evade host immunity and tolerate structurally diverse antibiotics.

## Methods

### Growth conditions and strains

*Mtb* strains (H37Rv genetic background) were cultured in a modified Sauton’s minimal medium: 0.5 g/L potassium phosphate dibasic, 0.5 g/L magnesium sulfate heptahydrate, 2.0 g/L citric acid monohydrate, and 0.05 g/L ferric ammonium citrate, supplemented with 0.05% (vol/vol) tyloxapol. The medium was supplemented with carbon sources used either individually or in combination, as either 0.1 mM cholesterol, 21.72 mM (2.0 g/L) glycerol, 5 mM acetate, 10 mM propionate, or other defined substrates as detailed in **Table S3**. For experimental conditions utilizing combinations of carbon sources (e.g., cholesterol + acetate), each substrate was maintained at the exact individual concentration specified in **Table S3**. Solid Middlebrook 7H10+OADC medium was used for transformation and CFU outgrowth. For the CRISPR interference (CRISPRi) KD strains of Mtb, we used the method described by Rock et al 2017^112^ to knock down the expression of genes of interest in Mtb and characterized dose response using MIC tests in the presence of appropriate antibiotics. For the overexpression strains, pDNTF and pDTCF expression vectors were used as described by Melief et al 2018.^113^ The list of strains generated in this study are listed in **Table S4**. When necessary, antibiotics were added to cultures (final concentrations): hygromycin - 50 µg/mL, kanamycin - 25 µg/mL, and anhydrotetracycline (ATc) - 100 ng/ml.

### Minimal inhibitory concentration (MIC)

For MIC assays, strains were grown in the Middlebrook 7H9+OADC medium until the mid-exponential phase (OD600nm ∼1) and washed twice in PBS buffer with 0.05% tyloxapol (PBST) and resuspended in a fresh modified Sauton’s minimal medium as single bacterial suspensions. The CRISPRi-mediated gene knockdown strain and the control non-targeting (NT) sgRNA strain of Mtb were pre-depleted in the presence of ATc for 5 d before assay for IC50 analysis. Cultures were then diluted back to OD600 of 0.05 and 80 μl cell suspension was plated in technical triplicate in wells containing the test compound and fresh ATc. For overexpression strains, the uninduced recombinants (-ATc) were used as control in comparison with overexpression (+ATc) strains. Drug compounds were solubilized in DMSO (or H2O + 0.3% tyloxapol for Vancomycin) and dispensed into 384-well plates using a Tecan’s D300e Digital Dispenser (HP). DMSO at a final concentration of 1% was used as no-drug control. In all, 80 μL of single-cell suspension was pipetted to each well and cultures were incubated for 14 days at 37 °C with 5% CO_2_. The plates were incubated for 14 days, and OD600nm was then recorded using a Tecan plate reader at days 0, 3, 7, 10 and 14. Results were represented as a percentage of 14-days area under the growth curve (AUC) of drug over no-drug control or the growth on lowest concentration where there was hormesis^114^ whichever was deemed appropriate according to the conditions. For all MIC tests, bacteria were seeded in 384-well plates and drugs were always dispensed by a D300e Digital Dispenser (HP), with DMSO being normalized to 1% (vol/vol) across wells. For all dose response curves, data represent the mean +/- s.d. for technical triplicates. Data are representative of at least two independent experiments. The IC50 values were derived using the four-parameter non-linear regression fit described in the data handling section and the dosing schemes were scaled around the expected IC50. For compounds lacking standalone activity (2MQ, pyridaben, and rotenone), dosing schemes were scaled from reported baseline concentrations of 1.5 µg/mL^46^, 12.5 µg/mL^46^, and 6.3 µg/mL^44^, respectively. OD600 measurements were used instead of colony-forming unit (CFU) enumeration to assess growth, as Rv1625c agonists exhibit minimal baseline bacteriostatic activity^106^, making CFUs unreliable for monitoring drug effects.

### DiaMOND assay to evaluate synergy of drug combinations

To evaluate the predicted synergy between selected compounds, we performed the diagonal measurement of n-way drug interactions (DiaMOND)^96,97^ assay as described by Cokol-Cakmak et al^96^. Bacterial culture inoculums were prepared as described in the MIC assays and seeded into 384-well plates. We first established single-drug IC50 values. Using these baseline values, we then determined the IC50 values for two- and three-drug combinations administered in fixed concentration ratios. The combination IC50 values were derived using the four-parameter non-linear regression fit described in the data handling section, with fixed-ratio dosing schemes scaled around the expected additive IC50 at 0.5x for two-drug combinations (tested from 0x to 1.0x in 0.1x increments of each drug) and 0.333x for three-drug combinations (tested from 0x to 0.667x in ∼0.067x increments of each drug) in such a way that their sum becomes 1x (e.g., in a two-drug combination, 0.5x of drug 1 and 0.5x of drug 2) as the mid-point. As noted previously, for compounds lacking standalone activity (2MQ, pyridaben, and rotenone), dosing schemes were scaled from reported baseline concentrations of 1.5 µg/mL^46^, 12.5 µg/mL^46^, and 6.3 µg/mL^44^, respectively, with the values for 2MQ and pyridaben corresponding to 5x their reported MICs^46^. Finally, the fractional inhibitory concentration (FIC2) was calculated to quantify the degree of interaction for each combination.

### Metabolic model refinement and curation

The iEK1011^71^ metabolic model of *Mycobacterium tuberculosis* H37Rv (Mtb) was updated with recent characterization of key metabolic processes, mass balanced reactions and newly characterized reactions to reflect the current Mtb metabolism. The major refinements include the reversibility of methylcitrate cycle (MCC)^36,72^ - changing the unidirectional reactions (2MCD, 2MCS, 2MID, 2MIL) into bi-directional reactions to allow the optimal use of reverse MCC in the presence of glycerol and glucose, whereas the MCC runs forward when in cholesterol+acetate^72^. The curation also included reactions that are related to the electron transport chain and electron transfer flavoproteins. The cytochrome BD has been characterized as non-proton pumping in Mtb^75–77^, hence we updated the reactions CYTBD and CYTBD2, and also balanced the proton-pumping in the electron-donating NADH dehydrogenases^44–46,78,79^. There were 3 reactions (NADH10, NADH2r, and NADH9) that represent Type I NADH dehydrogenases Nuo complexes. These reactions were mass balanced to include proton pumping. Based on the recent findings related to electron transfer flavoprotein characterization in Mtb^80^, the reactions ETF and ETFD were added as new reactions where EtfAB (Rv3028c and Rv3039c) re-oxidizes the FADH_2_ and transfers the electrons to EtfD (Rv0338c), which then reduces the electron carrier ubiquinone. This contributed to the removal of FADH_2_ which was replaced by ubiquinone as electron carriers for FRD and SUCDi reactions. We also added two new reactions for Succinate dehydrogenases to involve menaquinone as electron carrier^40,115^. Lactate dehydrogenase that uses NADH as cofactor (reaction: LDH_L) was removed and the quinone-dependent reactions (L_LACDcm, L_LACD, L_LACD2, L_LACD3) were updated to be encoded by Rv1872c (LldD2) and the Rv0694 (LldD1) was removed from the gene-protein-relationship (GPR) based on ^37,73^. The reaction, FPRA was mass balanced for the accurate electron representation for fdxrd[c] and fdxox[c] metabolites i.e, reduced-ferredoxin and oxidized-ferredoxin. The GPR for the reaction PPDK was removed based on the characterization that Rv1127c is no longer associated with this activity^72,74^. The inappropriate representation of acetate transport (Act2r) as antiport is now updated to be a symport with proton. Altogether 39 reactions were curated along with 54 reactions checked for consistency, 3 reactions were removed, 5 new reactions added and 8 new gene-protein-reaction associations were updated. The curated model, iSI1012, now has 1,012 genes, 1,232 reactions and 974 metabolites (**Table 1** and **Data S1**).

### Gene expression profiling following drug treatment

Wildtype Mtb was cultured in standard 7H9-rich media, then diluted back to OD600 = 0.1 in specific carbon source based modified Sauton’s media containing either DMSO as negative control, or the drug (GSK286) at specific concentration. Samples, in biological triplicates, were collected at specific timepoints. Samples were centrifuged at high speed for 5 min, supernatant was discarded and the cell pellet was immediately flash frozen in liquid nitrogen. Cell pellets were stored at -80°C until RNA extraction was performed as previously described.^116^

### Processing and analysis of RNA-Seq data

Sample collection and RNA-extraction was performed as described above. All samples were prepared with Illumina Stranded Total RNA Prep, Ligation with Ribo-Zero Plus (20040529 - Illumina, San Diego, CA). Total RNA samples were depleted of ribosomal RNA using the Ribo-Zero Plus rRNA Depletion Kit (Illumina, San Diego, CA). Quality and purity of mRNA samples was determined with 2100 Bioanalyzer (Agilent, Santa Clara, CA). All samples were sequenced on the NextSeq sequencing instrument (Illumina NextSeq 2000) using NextSeq 1000/2000 P2 Reagents (P2 XLEAP-SBS) flow cell v3. PhiX controls (Illumina, San Diego, CA) were used as a calibration control library for Illumina sequencing runs. Each sample library had 1% PhiX added and 750pM of the library was loaded onto the flow cells. Paired-end 150 bp reads were checked for technical artifacts using FastQC following Illumina default quality filtering steps. Raw FASTQ read data were processed using the R package DuffyNGS^117^ version 4. Briefly, raw reads were passed through a 2-stage alignment pipeline: (i) a pre-alignment stage to filter out unwanted transcripts, such as rRNA; and (ii) a main genomic alignment stage against the genome of interest. Reads were aligned to *M. tuberculosis* H37Rv (ASM19595v2) with Bowtie2^118^, using the command line option “very-sensitive”. BAM files from stage (ii) were converted into read depth wiggle tracks that recorded both uniquely mapped and multiply mapped reads to each of the forward and reverse strands of the genome(s) at single-nucleotide resolution. Gene transcript abundance was then measured by summing total reads landing inside annotated gene boundaries, expressed as raw read counts. We used these raw read counts as input for DESeq2^119^ for differential expression analysis and also converted the batch-wise reads into Transcripts Per Million (TPM) from raw read counts as the normalization for comparison across batches as well as for GraphPad Prism analyses. The RNA-Seq data of Mtb response to drug exposure generated for this study are available at the Gene Expression Omnibus under the accession number NCBI GEO: **GSE346561**.

### Incorporating drug treatment gene expression data on metabolic model

The iSI1012 metabolic model of Mtb was used for all the predictions in this study. For drug- and condition-specific models, we applied the gene expression data from both drug-treated and untreated control experiments using the GIMME^83^ algorithm on the iSI1012 model. This step was carried out to constrain the model to the specific condition being tested. We used GIMME because of the flexibility in defining objective function during implementation. The GIMME algorithm is implemented in the MATLAB_R2023b platform, using the “GIMME.m function” in the COBRA Toolbox after processing the gene expression data through “mapExpressionToReactions.m” function to convert the gene expression values as inputs to GIMME. The GitHub repository https://github.com/baliga-lab/iSI1012-metabolic-model-of-Mtb-and-Rv1625c-agonists-characterization contains all the condition-specific models generated in this study.

### Lipid extraction and thin-layer chromatography for PDIM estimation

Starter cultures of *Mycobacterium bovis* BCG Pasteur were grown in 7H9 broth with 10% OADC+glycerol and 0.05% Tyloxapol at 37°C until an A600 of 0.8. The grown cultures were centrifuged and pellets washed 3X times with 7H9 broth + 0.05% Tyloxapol before resuspending in the same media. The suspension was used as an inoculum to achieve an A600 of 0.1 in 7H9 + carbon source (glycerol, cholesterol or cholesterol+acetate). The cultures were incubated at 37°C until a A600 of 0.4, following which drug TBD11 or GSK286 were added. The cultures were incubated for a further 72h following which they were incubated with ^14^[C] acetic acid (10 µCi/ml) for 24h and then were harvested by centrifugation at 3,000 x g, 10 min, and washed twice with equal volumes of PBS. Wet weights of the pellets were recorded prior to proceeding with apolar lipid extraction as described by Dobson et al (1985)^120^. Apolar lipid extracts were resuspended in CHCl3:CH3OH (2:1 v:v) using volumes normalized to the original wet cell weight used for extraction. Equivalent volumes of sample were loaded on TLC and separated using petroleum ether: diethyl ether (9:1 v/v). Apolar lipids, including PDIMs were visualized by autoradiography using X-ray films exposed to the TLC plates for 3 days. A PDIM standard was run alongside the samples to confirm the position of PDIMs in all samples. Images were analyzed using ImageJ^121^ software for quantification of PDIM band intensity. The source file of images are available as extended data files.

### Data handling and analysis

All metabolic model simulations related to FBA were performed on the MATLAB_R2023b platform using the recent version of COBRA -The COnstraint-Based Reconstruction and Analysis toolbox (version 3.0)^122^. All condition-specific models were generated using GIMME^83^ algorithm. *In silico* flux predictions were performed using the in-built COBRA toolbox^122^ “optimizeCbModel”, “robustnessAnalysis” and “fluxVariability” function in MATLAB. The GitHub repository https://github.com/baliga-lab/iSI1012-metabolic-model-of-Mtb-and-Rv1625c-agonists-characterization contains all the details of the metabolic models and codes. **Data S1** contains detailed spreadsheets related to iSI1012 model. Experimental results were evaluated using GraphPad Prism version 11.0.2. All MIC graphs were generated by processing the data using R, followed by GraphPad Prism (non-linear regression fit of triplicates using log(inhibitor) vs response - variable slope (four parameters)) and assembled as composite figures in adobe illustrator version 30.4. The pathway illustration and ETC structures were drafted using adobe illustrator version 30.4.

### Quantification and statistical analysis

Statistical analyses reported in this article were performed using GraphPad Prism (version 11.0.2), R, or Python, with exact p-values method detailed in the corresponding figure legends and main text. Statistically non-significant (NS) analysis results were considered with p-value > 0.05 and other qualifying p-values were indicated accordingly ∗<0.05, ∗∗<0.01, ∗∗∗<0.001, and ∗∗∗∗<0.0001. To evaluate the statistical significance of the overlap between differentially expressed genes (DEGs) following GSK286 treatment and CRP target genes, a one-tailed hypergeometric test was conducted in R using the “phyper” function. Where applicable, resulting p-values were corrected for multiple hypothesis testing using the Benjamini-Hochberg false discovery rate procedure.

## Materials availability

This study did not generate new unique reagents.

## Data availability

The GitHub repository https://github.com/baliga-lab/iSI1012-metabolic-model-of-Mtb-and-Rv1625c-agonists-characterization contains all the GIMME derived condition-specific models generated in this study. All details of model curation are provided as supplementary materials. The RNA-Seq data generated for this study are available in the Gene Expression Omnibus under the accession number NCBI GEO: **GSE346561**.

## Code availability

R and MATLAB code, with data and description for implementation, is available in the GitHub repository https://github.com/baliga-lab/iSI1012-metabolic-model-of-Mtb-and-Rv1625c-agonists-characterization.

## Resource availability

**Lead contact** - Further information and requests for resources, strains and reagents should be directed to and will be fulfilled by the lead contact: Nitin Baliga.

## Supplementary Data

**Data S1:** Description of the iSI1012 metabolic model used in this study.

**Data S2:** Flux values from contextualized models and a comprehensive list of acetylated and succinylated enzymes.

**Data S3:** NAD(P)H-utilizing enzymes. Genes upregulated under conditions where GSK286 is inactive (i.e., cholesterol + acetate) are enriched for NAD(P)H-utilizing enzymes, which may help dissipate reductive stress during GSK286 treatment.

**Data S4:** Gene expression data related to heatmaps

## Supplementary Tables

**Table S1:** Model predicted inhibition versus non-inhibition by various substrates.

**Table S2:** IC50 values across all tested substrates.

**Table S3:** List of all substrates used in this study and their specific concentrations.

**Table S4:** Bacterial strains and materials used in this study.

## Supplementary Figures

**Figure S1:** Growth inhibition profiles in the presence of diverse carbon sources.

**Figure S2:** Correlation between model predicted NADH versus observed IC50.

**Figure S3:** DiaMOND assay results for the combinations.

**Figure S4:** Correlation between molecular weight of the compounds versus FIC.

## Source files

**Source 1:** Raw images of thin layer chromatography results

## Acknowledgments

We gratefully acknowledge Olalla Sanz, Elena Jimenez, Khisi Mdluli, Linu John, Helena Boshoff, Valerie Mizrahi, Dirk Schnappinger, and Tanya Parish for their insightful comments and feedback on this study. We gratefully acknowledge Tanya Parish, Renee Allen and Lauren Ames for their kind gift of pDTCF and pDTNF plasmids for overexpression strains. We also thank Betsy Russell (Bill & Melinda Gates Foundation) for sharing the TBD11 (mCLB073) compound used in this study. We are grateful to Helena Boshoff and Laura Cleghorn for their kind gift of 2MQ (2-Mercapto-Quinazolinones - Compound 7 (DDD00853663): N-(4,4-Difluorocyclohexyl)-2-((4-oxo-3,4-dihydroquinazolin-2-yl)thio)acetamide) from the University of Dundee, UK. We gratefully acknowledge Brittany Spanier-Marson and Aarun Hendrickson for their valuable technical support in compiling metadata information for the gene expression datasets. We thank Pamela Troisch and the Molecular & Cell Core Facility of the Institute for Systems Biology for their help with RNA sequencing. We thank members of the Baliga lab for critical discussions and feedback. Funding was provided by the National Institute of Allergy and Infectious Diseases of the National Institutes of Health (R01AI128215, R01AI141953, U19AI135976) and the Bill & Melinda Gates Foundation (INV-009322, INV-056403).

## Author contributions

Conceptualization: S.R.C.I., E.J.R.P., and N.S.B; Methodology: N.S.B., S.R.C.I., E.J.R.P., A.M.B. and K.A.H.; Investigation: S.R.C.I., J.D., A.K., M.P., and A.S.; Software: S.R.C.I., and W.-J.W.; Data curation: S.R.C.I., W.-J.W.; Formal analysis: S.R.C.I., K.A.H., and E.J.R.P.; Writing - Original draft, Review & Editing: S.R.C.I. and N.S.B.; Visualization: S.R.C.I. and N.S.B.; Resources: N.S.B. and A.M.B.; Supervision, project administration and funding acquisition: N.S.B.

## Declaration of interests

The authors declare no competing interests.

## Notes

https://www.ncbi.nlm.nih.gov/geo/query/acc.cgi?acc=GSE346561

https://github.com/baliga-lab/iSI1012-metabolic-model-of-Mtb-and-Rv1625c-agonists-characterization

### Competing Interest Statement

The authors have declared no competing interest.

