## Supplementary material for "An adenylyl cyclase switch reroutes carbon from growth to virulence lipids in *Mycobacterium tuberculosis*": Document S1

### Supplementary data information:

**Data S1:** Description of the iSI1012 metabolic model used in this study.

**Data S2:** Flux values from contextualized models and a comprehensive list of acetylated and succinylated enzymes.

**Data S3:** NAD(P)H-utilizing enzymes. Genes upregulated under conditions where GSK286 is inactive (i.e., cholesterol + acetate) are enriched for NAD(P)H-utilizing enzymes, which may help dissipate reductive stress during GSK286 treatment.

**Data S4:** Gene expression data related to heatmaps

**Supplementary Tables:**

**Table S1:** Model predicted inhibition versus non-inhibition by various substrates.

**Table S2:** IC50 values across all tested substrates.

**Table S3:** List of all substrates used in this study and their specific concentrations.

**Table S4:** Bacterial strains and materials used in this study.

**Supplementary Figures:**

**Figure S1:** Growth inhibition profiles in the presence of diverse carbon sources.

**Figure S2:** Correlation between model predicted NADH versus observed IC50.

**Figure S3:** DiaMOND assay results for the combinations.

**Figure S4:** Correlation between molecular weight of the compounds versus FIC.

**Source files:**

**Source 1:** Raw images of thin layer chromatography results

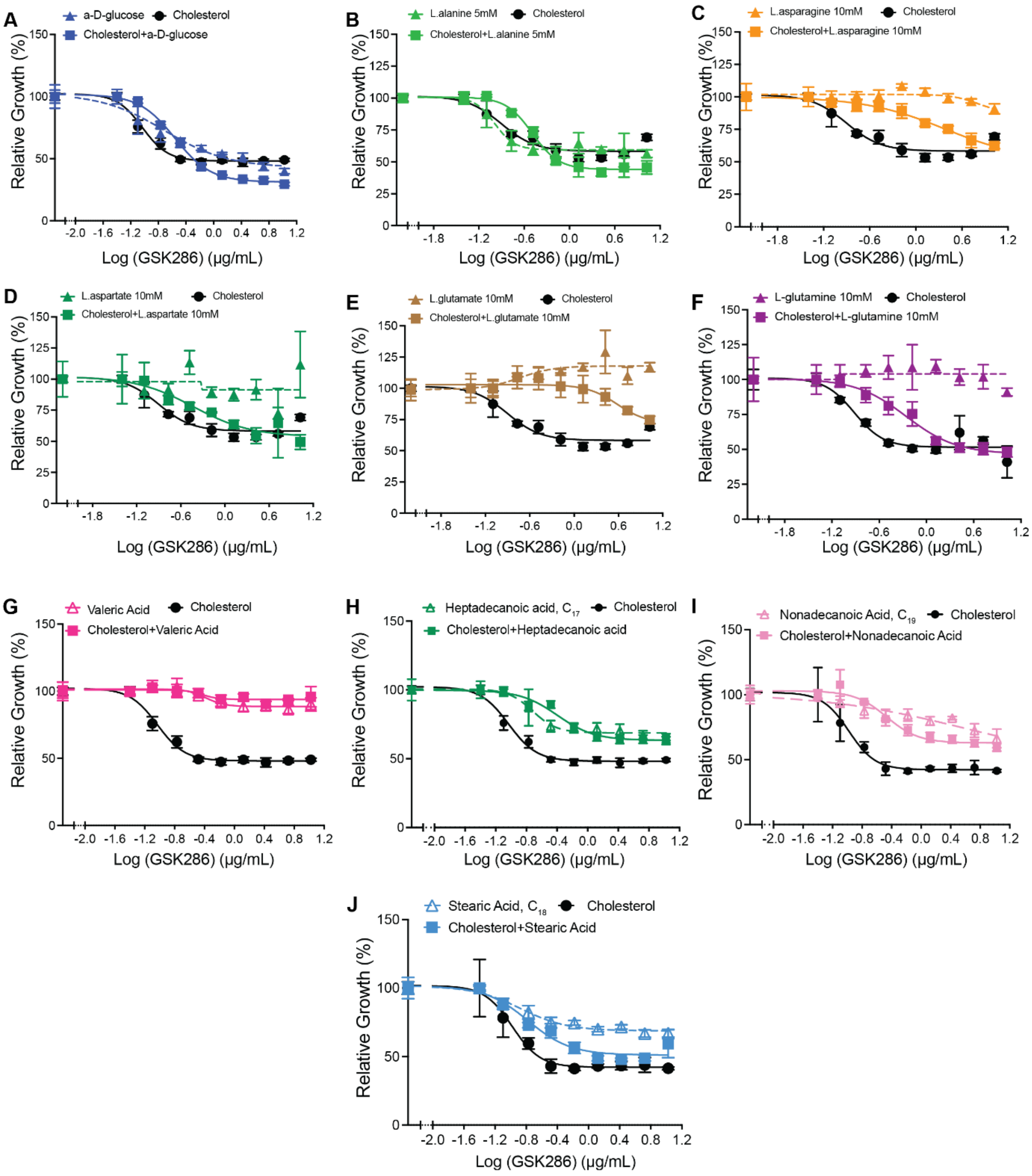


**Figure S1: Growth inhibition profiles in the presence of diverse carbon sources. A.** Glucose, **B.** L-alanine, **C.** L-asparagine, **D.** L-aspartic acid, **E.** L-glutamate, **F.** L-glutamine, **G.** Valeric acid, **H.** Heptadecanoic acid, **I.** Nonadecanoic acid, and **J.** Stearic acid. Data are representative of at least two independent experiments. Error bars correspond to standard deviation of replicates within the same experiment.

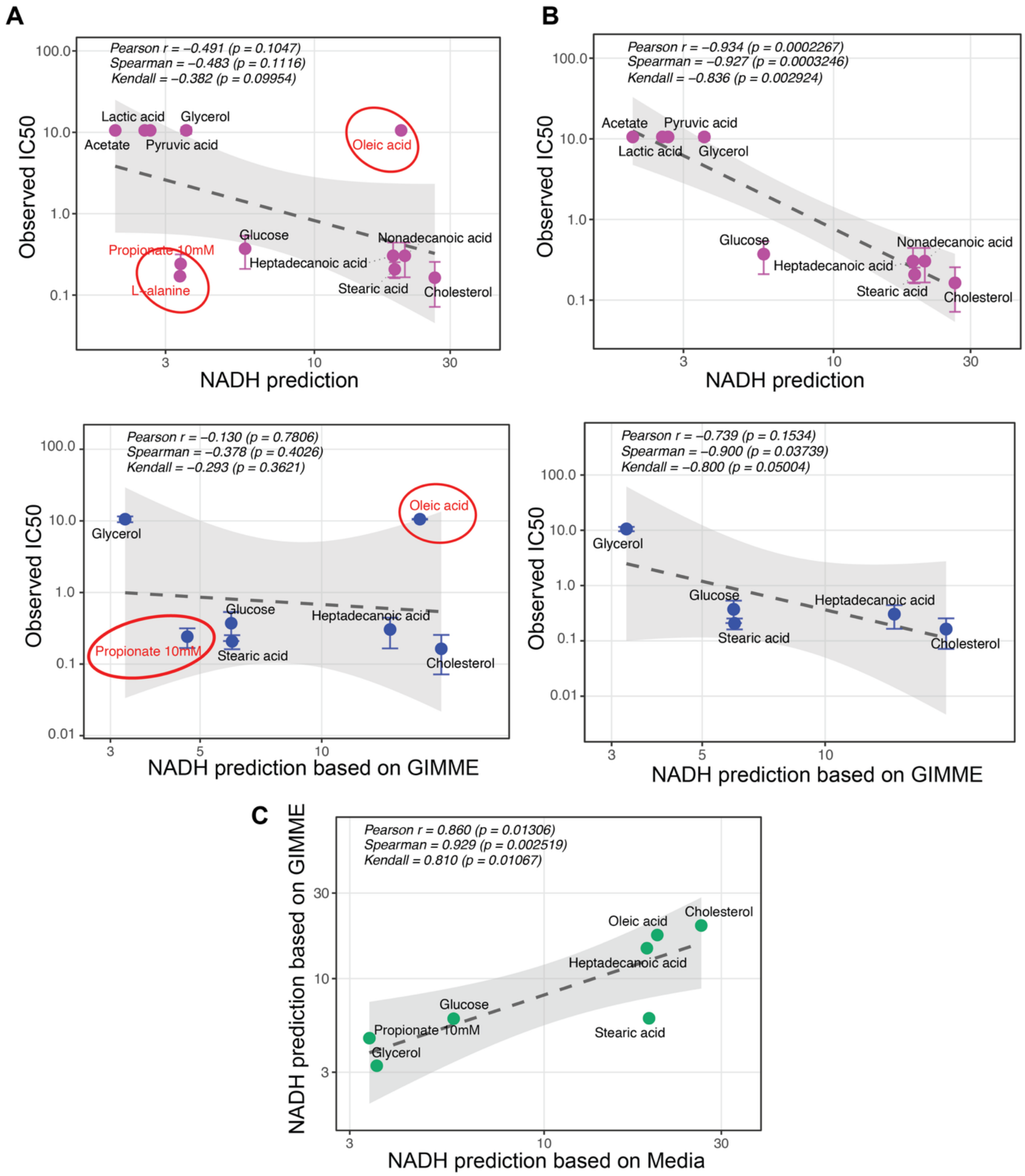


**Figure S2: Correlation between model-predicted NADH versus observed IC$_{50}$.** NADH levels were predicted using the media-constrained metabolic model as well as GIMME-based models by including a demand reaction to pool all the intracellular NADH. The plots shows before (**A**) and after (**B**) removing the outliers (circled in red). **C.** Correlation between media-constrained versus GIMME model NADH predictions.

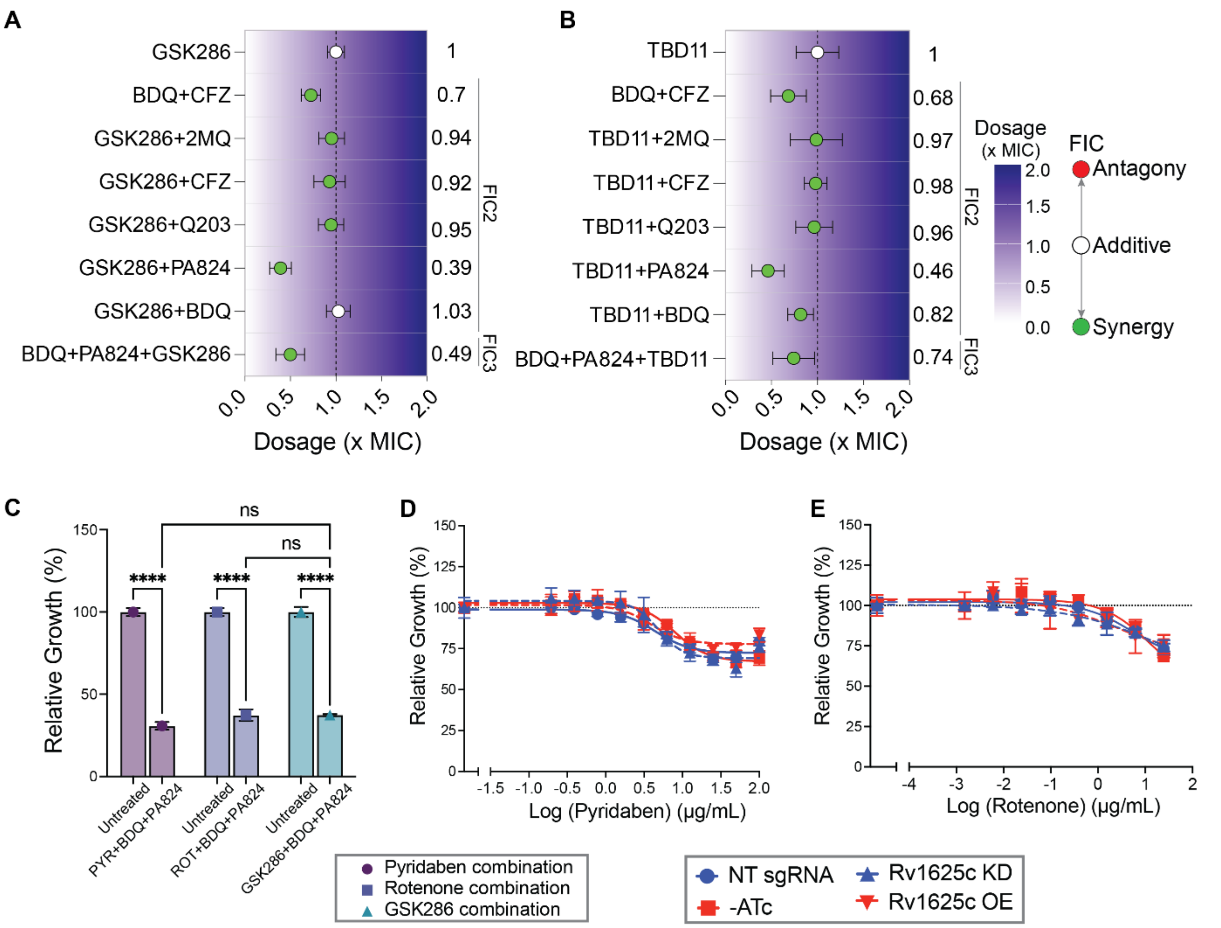


**Figure S3: DiaMOND assay results for the combinations. A.** GSK286 and **B.** TBD11 are synergistic with most of the ETC inhibitors with FIC ranging from mild to strong synergy. **C.** Pyridaben and rotenone share a similar synergy profile as GSK286 in the triple drug combinations. The activity of pyridaben (**D**) and rotenone (**E**) are tested using knockdown (blue) and overexpression (red) strains of Rv1625c and shows that these drugs are independent of Rv1625c.

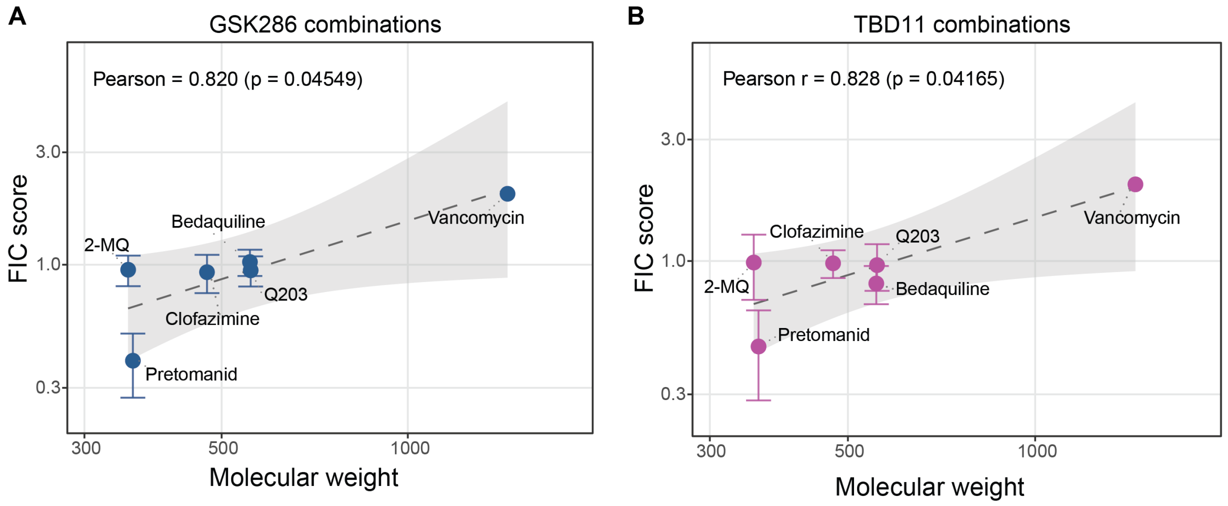


**Figure S4.** Correlation of molecular weight of the drugs tested in combination with GSK286 (**A**) and TBD11 (**B**) versus the fractional inhibitory concentration (FIC) scores from the diagonal assay. The higher the FIC, the drugs are antagonistic in the combinations.

**Table S1.** Predicted carbon sources that could rescue Rv1625c agonists activity in the presence of cholesterol

| Substrate | Exchange Reaction | Prediction | Observed* |
|---|---|---|---|
| Glycerol | EX_glyc | Non inhibitory | Non inhibitory |
| Acetate | EX_ac | Non inhibitory | Non inhibitory |
| L-Lactic Acid | EX_lac_L | Non inhibitory | Non inhibitory |
| Pyruvic Acid | EX_pyr | Non inhibitory | Non inhibitory |
| Citric Acid | EX_cit | Non inhibitory | |
| D-Galactose | EX_gal | Non inhibitory | |
| D-Serine | EX_ser_D | Non inhibitory | |
| Fumaric Acid | EX_fum | Non inhibitory | Not tested |
| a-Ketoglutaric Acid | EX_akg | Non inhibitory | |
| L-Proline | EX_pro_L | Non inhibitory | |
| L-Serine | EX_ser_L | Non inhibitory | |
| L-Threonine | EX_thr_L | Non inhibitory | |
| a-D-Glucose | EX_glc | Inhibitory | Inhibitory |
| Cholesterol | EX_chsterol | Inhibitory | Inhibitory |
| D-Alanine | EX_ala_D | Inhibitory | Inhibitory |
| Heptadecanoic Acid | EX_art_hepdcoa | Inhibitory | Inhibitory |
| L-Alanine | EX_ala_L | Inhibitory | Inhibitory |
| L-Glutamine | EX_gln_L | Inhibitory | Inhibitory |
| Nonadecanoic Acid | EX_art_nodcoa | Inhibitory | Inhibitory |
| Propionate 10mm | EX_ppa | Inhibitory | Inhibitory |
| Stearic Acid | EX_ocdca | Inhibitory | Inhibitory |
| L-Asparagine | EX_asn_L | Inhibitory | Inhibitory |
| L-Aspartic Acid | EX_asp_L | Inhibitory | Inhibitory |
| L-Glutamic Acid | EX_glu_L | Inhibitory | Inhibitory |
| Oleic Acid | EX_ocdcea | Inhibitory | Non-inhibitory |
| Valeric Acid | EX_art_ptcoa | Inhibitory | Non-inhibitory |
| Arachidic Acid | EX_art_arach | Inhibitory | |
| D-Ribose | EX_rib_D | Inhibitory | |
| D-Trehalose | EX_tre | Inhibitory | |
| Dl-Malic Acid | EX_mal_L | Inhibitory | |
| Formic Acid | EX_for | Inhibitory | |
| Glyoxylic Acid | EX_glyclt | Inhibitory | |
| L-Arabinose | EX_arab_L | Inhibitory | |
| Lauric Acid | EX_art_ddca | Inhibitory | Not tested |
| Maltose | EX_malt | Inhibitory | |
| N-Butyric Acid | EX_art_butcoa | Inhibitory | |
| Nonanoic Acid | EX_art_nndccoa | Inhibitory | |
| Palmitic Acid | EX_hdca | Inhibitory | |
| Pentadecanoic Acid | EX_art_pentdcoa | Inhibitory | |
| Phosphoenol Pyruvate | EX_art_pep | Inhibitory | |
| Succinic Acid | EX_succ | Inhibitory | |
| Heptanoic Acid | EX_art_hepcoa | Inhibitory | |

* The "non-inhibitory" class is defined based on >10-fold increase in $IC_{50}$ of GS286 in cholesterol

**Table S2:** IC50 values across all tested substrates.

| Substrate | Sole carbon (µg/ml) | In combination with cholesterol (µg/ml) |
|---|---|---|
| Cholesterol | 0.103 ± 0.018 | - |
| Glycerol | >10.56* | >10.56* |
| Other substrates tested | | |
| Propionate, C3 | 0.1189 ± 0.017 | 0.1821 ± 0.058 |
| L-alanine | 0.1706 ± 0.0038 | 0.3326 ± 0.13 |
| Heptadecanoic acid, C17 | 0.2668 ± 0.075 | 0.3403 ± 0.06 |
| a-D-Glucose | 0.3488 ± 0.31 | 0.3017 ± 0.04 |
| Stearic acid, C18 | 0.3732 ± 0.1 | 0.2243 ± 0.05 |
| Nonadecanoic acid, C19 | 1.849 ± 0.061 | 0.36 ± 0.10 |
| L-glutamine | 5.757 ± 0.67 | 0.34157 ± 0.08 |
| Acetate, C2 | >10.56* | >10.56* |
| Pyruvate | >10.56* | 0.6986 ± 0.11 |
| L-Lactate | >10.56* | >10.56* |
| L-asparagine | >10.56* | 1.2421 ± 0.19 |
| L-aspartate | >10.56* | 0.3960 ± 0.05 |
| L-glutamate | >10.56* | 2.723 ± 0.09 |
| Valeric acid, C5 | >10.56* | >10.56* |
| Oleic acid, C18:1 | >10.56* | >10.56 |
| Vitamin B12 | >10.56* | 0.2199 ± 0.041<br>>10.56* (+Propionate) |

*10.56 (µg/ml) was the upper limit of the MIC test - The MIC range evaluated in this study was capped at a maximum concentration of 10.56 µg/mL of GSK286.

**Table S3.** List of all substrates used in this study and their specific concentrations.

| Substrate | Class | Molecular weight | mM | g/L | CAS # | Sole Carbon | Cholesterol | Glycerol |
|---|---|---|---|---|---|---|---|---|
| Cholesterol | Sterols & Lipids | 386.65 | 0.1 | 0.0387 | 57-88-5 | + | NA | NA |
| Glycerol | Carbohydrate Intermediates | 92.09 | 21.72 | 2 | 56-81-5 | + | NA | NA |
| Other substrates tested | | | | | | | | |
| Acetate, C2 | Organic Acids / Short-Chain Fatty Acids | 82.03 | 5 | 0.41 | 127-09-03 | + | + | + |
| Propionate, C3 | Organic Acids / Short-Chain Fatty Acids | 96.061 | 10 | 0.961 | 137-40-6 | + | + | + |
| a-D-Glucose | Carbohydrates | 180.16 | 10 | 1.8 | 50-99-7 | + | + | + |
| Pyruvate | Carbohydrate Intermediates | 110.04 | 5 | 0.55 | 113-24-6 | + | + | + |
| L-Lactate | Organic Acids | 112.06 | 5 | 0.56 | 72-17-3 | + | + | + |
| L-alanine | Amino Acids | 89.09 | 5 | 0.45 | 56-41-7 | + | + | + |
| L-asparagine | Amino Acids | 132.12 | 10 | 1.32 | 70-47-3 | + | + | + |
| L-aspartate | Amino Acids | 192.12 | 10 | 1.92 | 3792-50-5 | + | + | + |
| L-glutamate | Amino Acids | 187.13 | 10 | 1.87 | 6106-04-03 | + | + | + |
| L-glutamine | Amino Acids | 146.14 | 10 | 1.46 | 56-85-9 | + | + | + |
| Valeric acid, C5 | Fatty Acids | 102.133 | 10 | 1.02133 | 109-52-4 | + | + | + |
| Heptadecanoic acid, C17 | Fatty Acids | 270.457 | 0.25 | 0.0676 | 506-12-7 | + | + | + |
| Stearic acid, C18 | Fatty Acids | 284.48 | 0.1 | 0.0284 | 57-11-4 | + | + | + |
| Oleic acid, C18:1 | Fatty Acids | 282.47 | 0.2 | 0.056 | 112-80-1 | + | + | + |
| Nonadecanoic acid, C19 | Fatty Acids | 298.51 | 0.1 | 0.029851 | 646-30-0 | + | + | + |
| Vitamin B12 | Vitamins & Cofactors | 1355.37 | 0.0074 | 0.01 | 68-19-9 | + | + | + |

**Table S4:** Bacterial strains and materials used in this study.

| Reagent/ Resource | Source | Identifier |
|---|---|---|
| **Bacterial strains** | | |
| H37Rv (*M. tuberculosis*) | D. Sherman lab | N/A |
| BCG (*M. bovis*) | D. Sherman lab | N/A |
| **Critical commercial assays** | | |
| NextSeq™ 1000/2000 P2 XLEAP-SBS™ Reagent Kit v3 | Illumina | 20100985 (300 cycles)<br>20100986 (200 cycles) |
| Illumina Stranded Total RNA Prep, Ligation with Ribo-Zero Plus | Illumina | 20040529 |
| RiboZero Plus rRNA Depletion kit | Illumina | 20036696 |
| RNA Prep Ligation | Illumina | 20040898 |
| cDNA Synthesis | Illumina | 20040896 |
| **Deposited data** | | |
| Raw and analyzed data | This paper | NCBI GEO: **GSE346561**<br>SRA Bio-project: PRJNA1526097 |
| GitHub Repository (Code and metabolic models) | This paper | GitHub: https://github.com/baliga-lab/iSI1012-metabolic-model-of-Mtb-and-Rv1625c-agonists-characterization |
| **Experimental models: Organisms/strains** | | |
| *M. tuberculosis*: CRISPRi knockdown of Rv1625c | This study | N/A |
| *M. tuberculosis*: CRISPRi knockdown of *icl1* | This study | N/A |
| *M. tuberculosis*: CRISPRi knockdown of *pckA* | This study | N/A |
| *M. tuberculosis*: Overexpression strain of Rv1625c | This study | N/A |
| **Recombinant DNAs and oligonucleotides** | | |
| CRISPRi sgRNAs | This study | N/A |
| Overexpression strain vectors (pDTCF and pDTNF) | Parish lab | N/A |
| **Software and algorithms** | | |
| DuffyNGS | Vignali et al. J. Clin. Invest., 121 (2011), pp. 1119-1129, 10.1172/JCI43457 | N/A |
| GraphPad Prism, Version 11.0.1 | GraphPad Software | https://www.graphpad.com/ |
| ImageJ | Schneider et al. Nat. Methods, 9 (2012), pp. 671-675, 10.1038/nmeth.2089 | https://imagej.net/ij/ |

**Source files**:

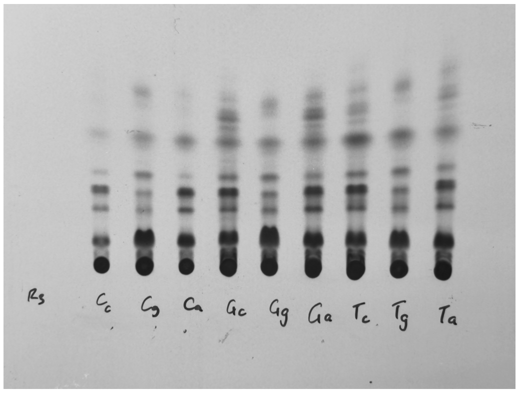

R3 - Replicate 3 (used in **Figure 4F**)

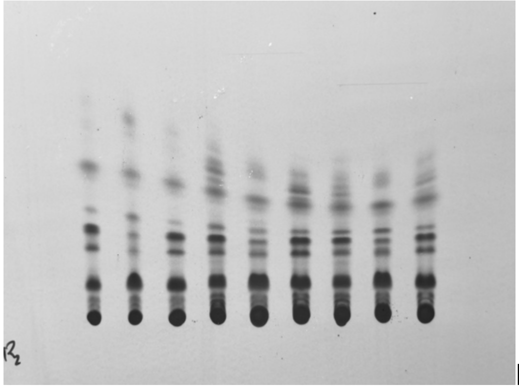

R2 - Replicate 2

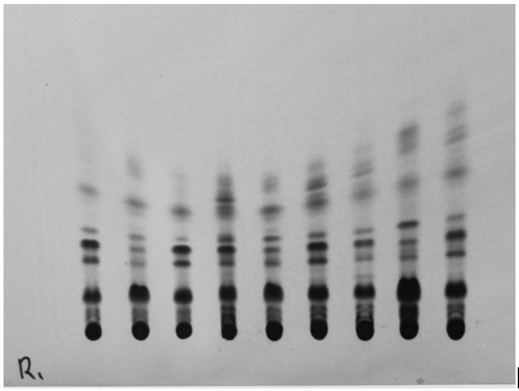

R1 - Replicate 1

**Source 1. Thin layer chromatography (TLC) images:** Replicate 3 inset labels correspond to sample identities: **Cc** – Control cholesterol; **Cg** – Control glycerol; **Ca** – Control cholesterol + acetate; **Gc** – GSK286 + cholesterol; **Gg** – GSK286 + glycerol; **Ga** – GSK286 + cholesterol + acetate; **Tc** – TBD11 + cholesterol; **Tg** – TBD11 + glycerol; **Ta** – TBD11 + cholesterol + acetate; Replicates 2 and 1 follow the same order of sample loadings. Replicate 3 image was used in **Figure 4F** and the intensities were measured from all three replicates using ImageJ for the **Figure 4G**.
